# Organoid-based human stomach micro-physiological system to recapitulate the dynamic mucosal defense mechanism

**DOI:** 10.1101/2022.03.02.482603

**Authors:** Hye-Jin Jeong, Ji-Hyeon Park, Joo H. Kang, Seong-Ho Kong, Tae-Eun Park

**Affiliations:** Department of Biomedical Engineering, Ulsan National Institute of Science and Technology, Ulsan 44919, South Korea; Department of Surgery, Seoul National University Hospital, Seoul National University College of Medicine, Seoul 03080, South Korea

## Abstract

Several stomach diseases are attributed to the dysregulation of physiological function of gastric mucosal barrier by pathogens. Gastric organoids are a promising tool to develop treatment strategies for gastric infections. However, their functional features of *in vivo* gastric mucosal barrier and host-microbe interactions are limited due to the lack of physiological stimuli. Herein, we describe the first human stomach micro-physiological system (hsMPS) with physiologically relevant gastric mucosal defense system based on the combination of organoid and MPS technology. A fluid flow enhanced epithelial-mesenchymal interaction in the hsMPS enables functional maturation of gastric epithelial cells, which allows for the recreation of mesh-like mucus layer containing high level of mucus protective peptides and well-developed epithelial junctional complexes. Furthermore, gastroprotection mechanisms against *Helicobacter pylori* are successfully demonstrated in our system. Therefore, hsMPS represents a new *in vitro* tool for research where gastric mucosal defense mechanism is pivotal for developing therapeutic strategies.

## Introduction

The human gastric mucosal barrier is composed of epithelial, mucus, and submucosal elements that play gastroprotective roles by blocking pathogens and harmful substances (*1*). The gastric epithelial barrier is maintained by continuous renewal of gastric stem cells within a stem cell niche where increased canonical wingless/integrated (Wnt) signaling activity is sustained. The surrounding mesenchymal stromal cells in the lamina muscularis mucosae influence the gastric epithelium function as key components of the gastric stem cell niche (*2*). The gastric stem cells differentiate into functional pit cells, which contribute to the overall function of the stomach, including secretion of mucus element for gastric protection and controlling the stomach's pH environment. The gastric epithelium is covered by a mucus layer, which acts as a protective viscoelastic barrier composed of highly glycosylated mucin proteins forming a net-like polymer as well as protective peptides including trefoil factor peptides (TFF1 and TFF2) (*3*). In the submucosal compartment, mucosal blood flow supplies the mucosa with nutrient and oxygen and removes metabolic waste for maintaining the mucosal barrier and provides circulating immune cells for gastric immune responses against pathogens.

The homeostatic disruption of the gastric mucosal defense system against various foreign agents passing through the stomach leads to various kinds of stomach disease pathogenesis. For example, *Helicobacter pylori* (*H. pylori*) infection is a leading cause of peptic ulcer disease, chronic gastritis, and stomach cancer, as it mediates a compromise of the gastric mucosal barrier affecting the integrity of stomach (*1*). Therefore, understanding the gastric mucosal defense mechanisms against pathogens, and how pathogens evade or exploit the physiological defense system of the host is important to develop effective treatment strategies against gastric diseases.

Three-dimensional (3D) gastric organoids derived from stomach tissue are a promising tool for treatment development against gastric infections and disorders (*4, 5*), since they contain self-renewing gastric stem cells that differentiate into multiple gastric-specific epithelial cell types embedded in an extracellular matrix (ECM) hydrogel (*6*). 3D gastric organoids are more physiologically relevant compared to stomach cell lines since they mimic physiological epithelial regeneration process (*6*) and *H. pylori* pathogenesis (*7*). However, the current gastric organoid systems have clear disadvantages in recreating the complex gastric mucosal defense system. In human gastric tissue, various niche factors in gastric glands, including gastric mesenchymal stromal cells (gMSCs) and extracellular matrix (ECM) help to maintain the orchestrated stem cell proliferation and differentiation to provide the functional epithelial cells constituting the gastric barrier (*8, 9*). In gastric organoid culture, however, proliferation and differentiation of the gastric stem cells are simply regulated by the addition or withdrawal of Wnt signaling molecules (Wnt3a and R-spondin1) in the culture media, which results in an unbalanced generation of gastric epithelial cells (*5*). In addition, cellular interaction between epithelial and immune cells, which induces inflammatory responses (*10*), is not reproduced in organoid cultures. The enclosed lumen structure of the gastric organoids also hampers access to the apical side of the gastric epithelium, as well as the excretion of metabolic wastes from pathogens and epithelial cells, which is an inaccurate representation of the cell responses to pathogens (*2*). Since gastric organoids have immature characteristics and with the technical difficulty associated with the model system, there remains a need to develop technologies that promote the maturation of gastric organoids with physiologically relevant features of gastric mucosal barrier and host-microbe interactions.

Recently, considerable efforts have been made to combine organoid technology and micro-physiological systems (MPSs) to develop a more powerful *in vitro* platform with a synergistic combination of their features (*11*). MPS is a microfabricated 3D cellular construct designed to recapitulate the functions of organs through engineered microenvironment control (*11*). Implantation of brain (*12*), intestinal (*13*), ocular (*14*), and kidney organoid (*15*) in MPS devices has led to greater functionality of organoids owing to controlled biochemical and biophysical microenvironmental cues and dynamic cell-cell interplay (*16*). An MPS system providing peristaltic flow through enclosed gastric organoids has been previously developed for efficient luminal delivery of nutrients or pharmacological agents (*16*). However, the implanted gastric organoid showed no physiological improvement, thereby demonstrating its biological and engineering design constraints.

Herein, we describe the first human stomach MPS (hsMPS) with a greatly enhanced gastric mucosal barrier based on the combination of MPS technology with organoids. The hsMPS where epithelial cells derived from human antral organoids (hAOs) and primary gMSCs extracted from stomach tissue are cultivated under controlled flow recapitulates gastric epithelial homeostasis for maintenance of the functional mucosal barrier. The resulting hsMPS exhibits physiologically relevant mucosal defense system, including mesh-like mucus layer, which continuously covers the gastric mucosa and contains TFF peptides, and enhanced gastric epithelial junctional complexes. Furthermore, *H. pylori* infection model was established in the hsMPS co-cultured with peripheral blood mononuclear cells (PBMCs) to monitor the dynamic mucosal defense mechanism against *H. pylori* infections.

## Results

### Reconstitution of the orchestrated gastric epithelial homeostasis in an MPS system

Balanced between gastric stem cell self-renewal and differentiation is a key for the formation of the gastric mucosal barrier, therefore, microenvironmental cues for gastric epithelial homeostasis were explored to generate the functionally enhanced gastric organoid-based model (**Fig. 1A**). The gMSCs located at the lamina muscularis mucosae directly beneath the gastric glands have been suggested to play a key role in maintaining stem cell activity and proliferation as an important gastric stem cell niche (*8*). We hypothesized that co-cultivation of gastric epithelial cells with gMSCs could create the gastric stem cell niche-like environment that provides paracrine signals to maintain epithelial stemness and enable continuous supply of gastric progenitors in differentiation media (DM) condition where differentiation progeny can be actively generated as well. Although it has been documented that gMSCs sustain the gastric stem cell activity as a key cellular component of stem niche, maintaining their capacity *in vitro* is critical in achieving our goal. We assumed that mimicking mucosal blood flow (*13, 17*) delivering the nutrients and eliminating waste would improve the functional maintenance of gMSC as well as gastric epithelium (*18*), and facilitate the epithelial-mesenchymal interactions.

To explore these possibilities, a compartmentalized polydimethylsiloxane (PDMS) device with luminal and abluminal microchannels separated by a porous polyethylene terephthalate (PET) membrane, which permits dynamic cell-cell crosstalk and fluid flow control, was used in this study (**Fig. 1B**). The human stomach was reconstituted via an MPS device by culturing singularized hAOs on the luminal (upper) microchannel interfaced with autologous primary gMSCs on the abluminal (lower) channel. The hAOs were particularly used to model the gastric antral mucosa where initial *H. pylori* colonization process occurs to study host-bacterial interactions at early stage. We first plated gMSCs in a lower microchannel and the device was flipped immediately to allow for gMSCs attachment to the bottom side of PET membrane. At 2 h following gMSCs seeding, gastric epithelial cells prepared by singularization of the hAOs were seeded on the apical channel and cultured for four days in static conditions in expansion media (EM). On day 5, Wnt protein-deprived DM was supplied to luminal and abluminal channels at a continuous flow at 60 μL h^−1^ (**Fig. 1C**) (*13, 17*). Confocal fluorescence microscopy was conducted on day 6, which revealed a distinct epithelial monolayer on the PET membrane in the luminal microchannel with well-developed junctional complexes which contained E-cadherin along the lateral borders (**Fig. 1D**). Gastric surface mucous cells secreting MUC5AC and TFF1 and deep mucous gland cells secreting MUC6 were successfully identified as well in the hsMPS (**Fig. 1D**). Furthermore, the gastrin-producing G cells, found only in the gastric antral region, were detected using immunostaining for gastrin (**Fig. 1D**), proving that multiple types of functional gastric epithelial cells were successfully generated in a hsMPS system.

**Fig. 1.**
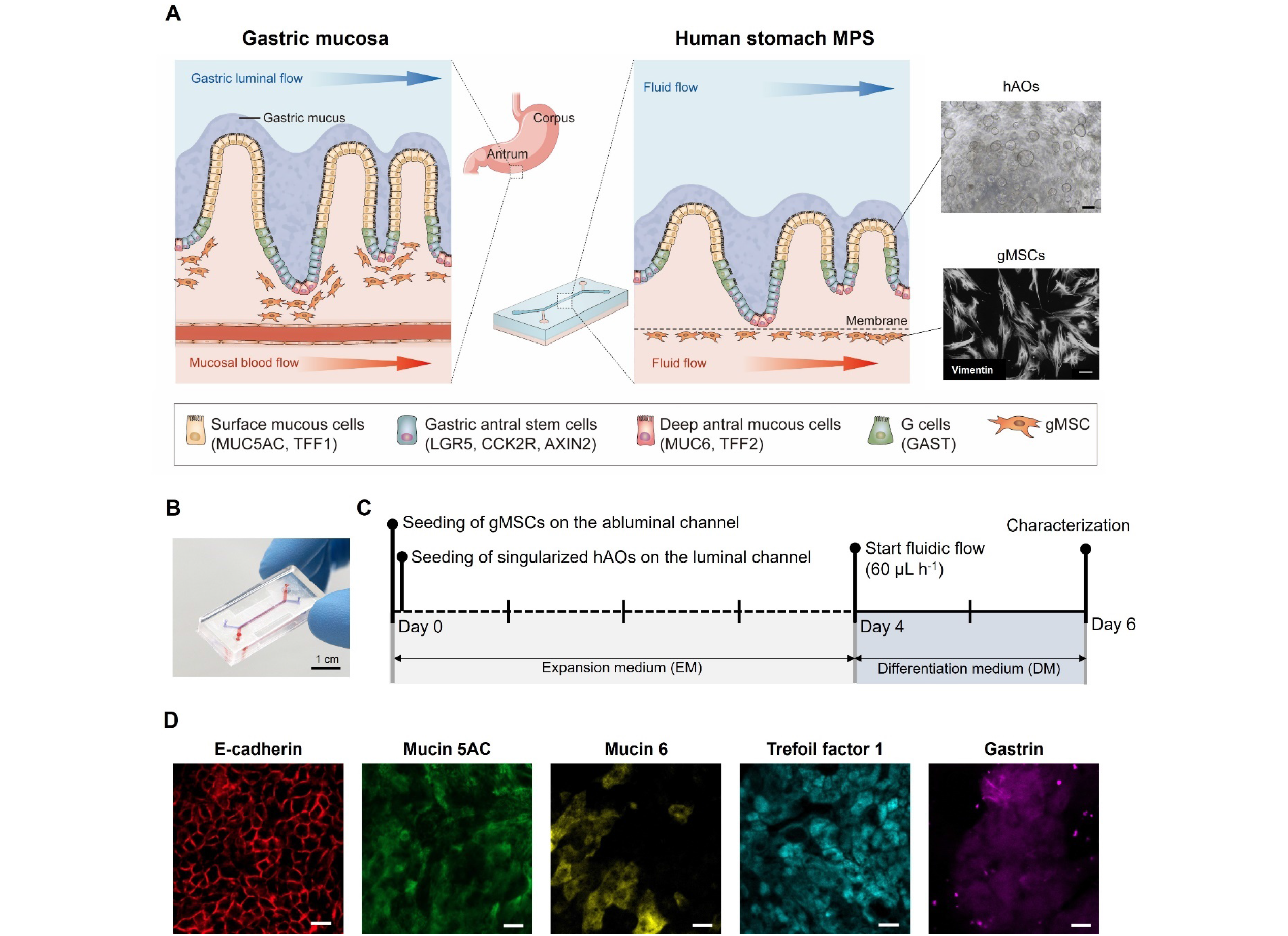
Reconstitution of the human stomach in an MPS device. (A) Schematic illustration of the human gastric mucosa (left) and the human stomach MPS (hsMPS) with singularized human gastric antral organoid (hAOs) cultured on the apical luminal channel interfaced with primary gastric stromal cells (gMSCs) on the basal abluminal channel (center). At the top right, a bright field image of hAOs (bar, 250 μm) used to generate gastric epithelium, and at the bottom right, an immunofluorescence micrograph of the gMSCs labelled with Vimentin (bar, 100 μm) are shown. Photograph of the hsMPS (bar, 1 cm). (C) Timeline for the reconstitution of the human stomach in an MPS device. (D) Immunofluorescence micrographs of the gastric epithelium in the hsMPS at 6 days labelled with E-cadherin, mucin 5AC, mucin 6, trefoil factor 1, and gastrin (bar, 20 μm).

### Fluid flow-enhanced niche function of gMSC to sustain gastric stem cell activity in the hsMPS

We determined whether co-culturing of gMSCs in the presence of fluid flow in an MPS device would enhance the maintenance of gastric epithelial stemness in DM. We assessed the mRNA expressions of genes encoding the major gastric stem cell markers LGR5, CCK2R, and AXIN2 in the hsMPS compared to hAOs and gastric antrum tissue (**Fig. 2A** and **fig. S1**). LGR5- and CCK2R-positive cells are long-lived stem cells that reside in the deep base of the antral glands responsible for maintaining gastric epithelial homeostasis (*3*). AXIN2 is also a known marker of gastric stem cells with high proliferation rate largely dependent on the Wnt pathway (*19*). We found that the relative mRNA expression of genes encoding LGR5 and CCK2R was significantly upregulated in co-culture compared to that in mono-culture conditions only in the presence of fluid flow (**Fig. 2A** and **fig. S1**), indicating that constant flow greatly influenced the niche function of gMSCs that sustains gastric stem cell activity. Importantly, under these conditions in the hsMPS, gastric epithelial cells exhibited similar mRNA expression level of CCK2R and AXIN2 with human gastric tissue, while conventional hAOs cultures show significantly different expression levels for all three stem cell markers compared to *in vivo* (**Fig. 2A**). These results highlight the synergistic effect of co-cultures with gMSCs and fluid flow in recapitulating physiological gastric stem cell homeostasis in the hsMPS.

**Fig. 2.**
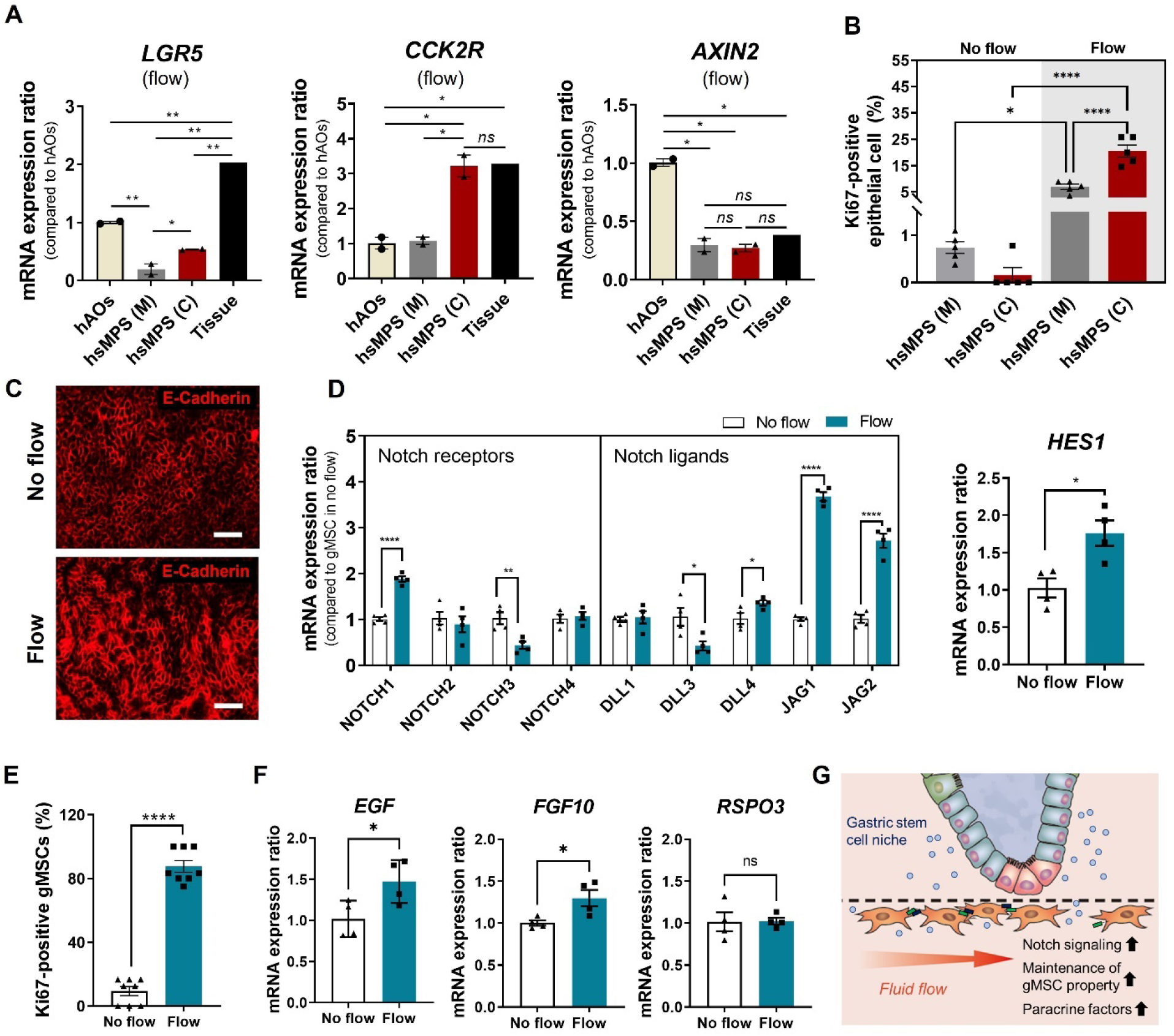
Gastric epithelial mitotic activity and morphogenesis in the hsMPS. (A) mRNA expression ratios of stem cell markers LGR5 (left), CCK2R (center), and AXIN2 (right) in gastric epithelial cells in the hsMPS in mono-culture (M) and co-cultures (C) under flow conditions compared to hAOs. (B) Percentage of Ki-67-positive epithelial cells analyzed using ImageJ under static and flow conditions. (C) Microscopic images (left; bar, 100 μm) and immunofluorescence confocal images (right; bar, 50 μm) of gastric epithelium co-cultured with gMSCs labelled with E-cadherin. (D) mRNA expression ratios of genes encoding Notch receptors (NOTCH1, NOTCH2, NOTCH3, and NOTCH4), Notch ligands (DLL1, DLL3, DLL4, JAG1, and JAG2), and Notch target HES1 in gMSCs in flow versus static conditions. (E) Percentage of Ki-67-positive gMSCs analyzed using ImageJ under static and flow conditions. (F) mRNA expression ratios of genes encoding paracrine factors—EGF (left), FGF10 (center), and RSPO3 (right) in gMSCs in MPS device in flow versus static conditions. (G) Schematic illustration of fluid flow-enhanced gMSC function as a cellular niche factor. The results are presented as the mean ± s.e.m. For statistical analysis, an unpaired t-test was performed (\**P* < 0.05; \*\**P* < 0.01; \*\*\**P* < 0.001; \*\*\*\**P* < 0.0001).

Next, we determined whether gMSCs and fluid flow facilitated self-renewal capacity of gastric epithelium in an MPS device, which is essential for constant supply of progenitor cells to maintain gastric barrier. Confocal microscopy of Ki-67 was conducted to examine the mitotic activity of epithelium in flow versus static conditions and in co-cultures versus mono-cultures (**Fig. 2B** and **fig. S2**). In both mono- and co-culture conditions, exposure to fluid flow revealed significantly higher levels of Ki-67 in gastric epithelial cells, indicating that fluid flow stimulated mitotic activity of gastric epithelial cells (**Fig. 2B**). In good agreement with the mRNA data of stem cell markers (**fig. S1**), only when exposed to fluid flow, a great increase in the percentage of Ki-67-positive epithelial cells was observed in co-cultures versus mono-cultures (20±5.6% versus 6.9±1.9%) (**Fig. 2B**). After confirming that fluid flow is required to induce gMSCs-mediated gastric epithelial proliferation, we determined if the gastric epithelial invagination could be recapitulated in the hsMPS under these conditions. The invagination is essential for the gastric epithelium formation in 3D tissue during morphogenesis (*20*); specifically, proliferation of epithelial cells is an important factor in generating the epithelial invagination in periodic boundary conditions by inducing apical constriction (*20*). Confocal microscopic analysis of E-cadherin in the hsMPS revealed greater epithelial invagination in flow conditions due to the greater fraction of dividing cells, while the epithelium remained flat without fluid flow (**Fig. 2C**). To investigate how fluid flow facilitates the niche function of gMSCs in our stomach system, Notch activation in gMSCs, which is an important regulator of MSC maintenance (*21*), was assessed when cultured within the device in the absence of gastric epithelial cells. The gMSCs remain in undifferentiated state at lamina muscularis mucosae functioning as a cellular niche factor (*3*), however it is challenging to maintain their characteristics *in vitro* (*21*). Interestingly, the flow-enhanced Notch signaling factors was observed by comparing the gene expression level of Notch ligands and receptors in gMSCs in flow versus static conditions. Statistically significant increases in mRNA expression were observed for genes encoding Notch ligands JAG1, JAG2, and DLL4, and Notch receptor NOTCH1 (Notch activators), on the contrary, Notch ligand DLL3, an inhibitory Notch ligand, was decreased (**Fig. 2D**). Accordingly, HES1, a main downstream Notch target gene that inhibits the cell senescence and maintains the stemness of gMSCs (*21*) was significantly upregulated (**Fig. 2D**). Using Ki-67 assessment, it was also confirmed that gMSCs maintained mitotic activity when exposed to fluid flow in an MPS device due to the enhanced Notch signaling, while those in static conditions lost proliferative capacity (**Fig. 2E** and **fig. S2**). Important paracrine factors for the antral stem cell niches such as fibroblast growth factor 10 (FGF10) and epithermal growth factor (EGF) (*3*) that regulate self-renewal of stem cells were found upregulated in gMSCs in flow conditions (**Fig. 2F**). On the other hand, mRNA expression of R-Spondin 3 that activates Axin2+/Lgr5- was unchanged (*19*) (**Fig. 2F**), which may explain why AXIN2 marker was not upregulated in co-cultures versus mono-cultures in the hsMPS (**Fig. 2A**). These data reveal how fluid flow enhances the function of mesenchymal component in facilitating gastric epithelial homeostasis in the hsMPS, which highlights the advantage of the use of microfluidic system (**Fig. 2G**).

### Formation of *in vivo* relevant mucosal barrier with gMSC co-culture in flow conditions

We initially explored microenvironmental cues to achieve orchestrated gastric epithelial homeostasis in the hsMPS that are key factor for recapitulating the formation and maintenance of gastric mucosal barrier. Since we demonstrated the contribution of fluid flow in the epithelial-mesenchymal interaction in the hsMPS, we focused on the role of gMSCs in the recreation of gastric mucosal defense system in an MPS device under flow conditions. We first observed the mRNA expression of genes encoding major gastric mucus components (**Figs. 3A-3D**). The highly glycosylated-gel-forming mucins of the stomach including MUC5AC mucin, expressed in superficial gastric epithelium, and MUC6 mucin, found in gastric glands, physically protect the underlying gastric epithelium against the noxious agents. TFF peptides (TFF1 and TFF2) co-expressed with mucin proteins function as a chemical barrier in gastric mucus layer interacting with pathogens bound to the mucins (*3*). The TFF1 offers binding moiety for *H. pylori* at the gastric antrum mucus layer and protects the gastric epithelium during *H. pylori* infection (*22, 23*); meanwhile, TFF2 plays a critical role in maintaining gastric mucosal integrity and exerts antibiotic function against pathogens entering the stomach (*24*). High similarity to the *in vivo* gastric mucous barrier in the hsMPSs was observed when mucin-related gene expression of gastric epithelial cells was compared with that of hAOs. We found statistically significant increases in mRNA expression of genes encoding MUC5AC, MUC6, TFF1, and TFF2 in the hsMPS compared to hAOs (**Figs. 3A-3D**). Furthermore, the mRNA levels of those genes were significantly upregulated and better matched with the *in vivo* gastric tissue in gastric epithelial cells when co-cultured with gMSCs compared to mono-culture in the hsMPS, as expected (**Figs. 3A-3D**). Interestingly, the mRNA expression of genes encoding TFF1 and TFF2 was improved in hsMPS in co-culture conditions, although their expression was still lower compared to *in vivo* tissues (**Figs. 3C and 3D**). Considering that the current *in vitro* gastric models, including hAOs and human stomach cell lines, express extremely low level of TFF peptides, which necessities the transfection-mediated TFF overexpression (*25*), hsMPS may provide a promising *in vitro* tool to study the gastric physiology associated to chemical barrier of the gastric mucus layer *in vitro*.

Confocal microscopic analysis confirmed that upregulated MUC5AC successfully formed the mesh-like mucin network on the gastric epithelium in co-culture conditions (**Fig. 3E**) in an MPS device, which is typically observed in adherent mucus layer in gastric tissue (*26*). The expression of MUC5AC was also confirmed in the absence of gMSCs, however, formation of the mucin network was not observed (**Fig. 3E**). These results show that gMSCs enhanced the dynamic balance of production, secretion, and polymerization of mucin in the MPS under fluid flow. A higher level of MUC6 mucin protein expression was also identified using immunofluorescence (**Fig. 3F**), which revealed that epithelial cells were efficiently differentiated in deep mucous gland cells as well.

**Fig. 3.**
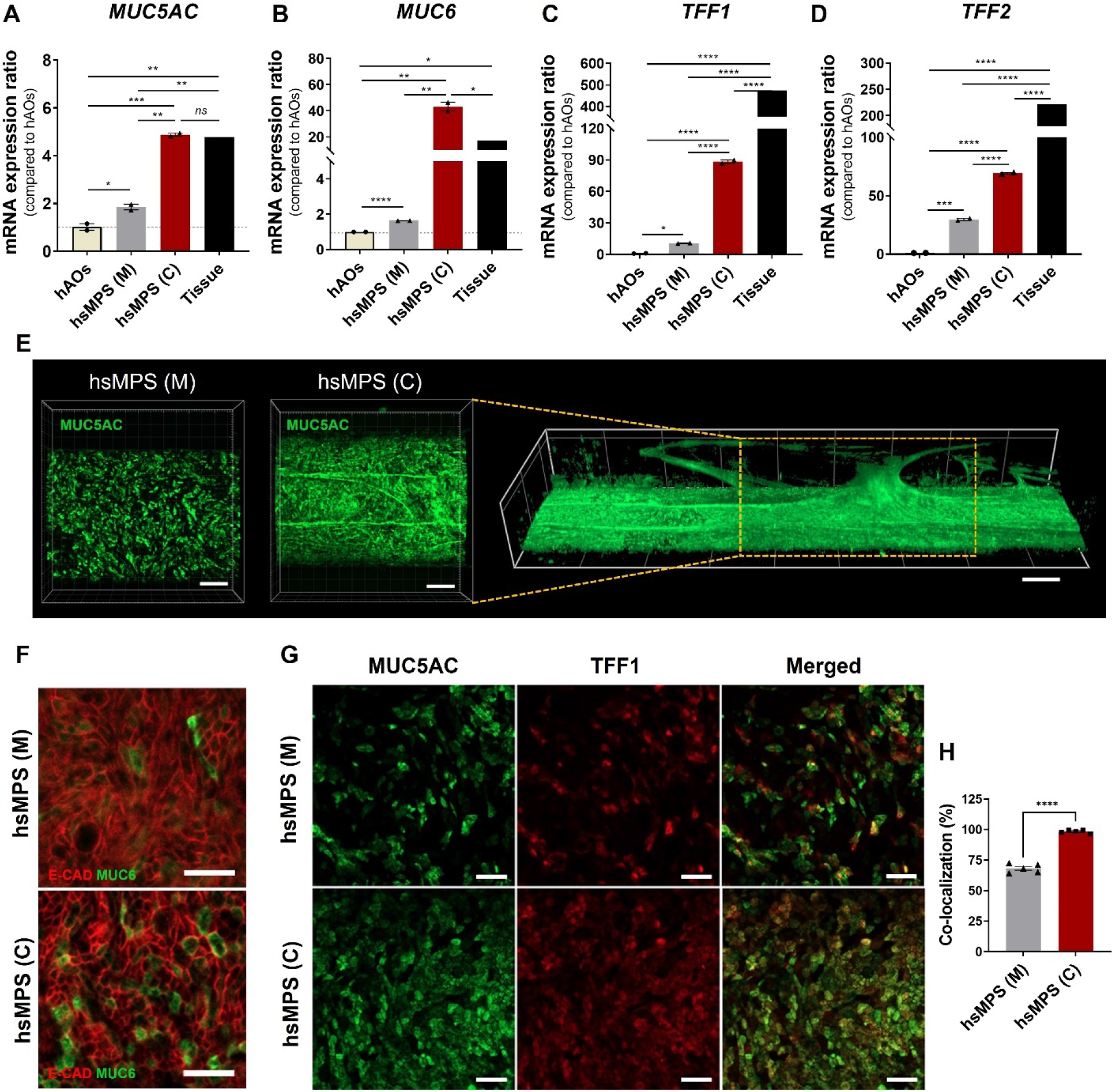
Recapitulation of *in vivo* relevant gastric mucous barrier in the hsMPS. (A-D) mRNA expression ratios of genes encoding mucin-associated markers MUC5AC (A), MUC6 (B), TFF1 and TFF2 (D) in the gastric epithelial cells in mono-cultures (M) and co-cultures (C) conditions in the hsMPS in the presence of fluid flow compared to hAOs. (E) Immunofluorescence micrograph of gastric epithelium labelled with mucin 5AC (green) under mono-culture (left) and co-culture (center and right) conditions (bar, 200 μm). (F) Immunofluorescence micrographs of epithelial cells stained with mucin 6 (green) and E-cadherin (red) in mono-culture (top) and co-culture (bottom) (bar, 50 μm). (G) Immunofluorescence images of gastric epithelial cells in monoculture and co-culture conditions labelled with mucin 5AC (green) and TFF1 (red) (bar, 50 μm). (H) Co-localization efficiency of TFF1 with MUC5AC analyzed using ImageJ. The results are presented as the mean ± s.e.m. For statistical analysis, an unpaired t-test was performed (\**P* < 0.05; \*\**P* < 0.01; \*\*\**P* < 0.001; \*\*\*\**P* < 0.0001).

We further identified the expression and localization patterns of MUC5AC and TFF1 in the hsMPS (**Fig. 3G**). Higher protein expression of MUC5AC and TFF1 was observed when co-cultured with gMSCs using confocal immunofluorescence, consistent with data from qRT–PCR. The co-localization efficiency of TFF1 with MUC5AC was highly enhanced in the presence of gMSCs (99.6% versus 68.1%), proving that our hsMPS successfully recapitulated the TFF1 binding to MUC5AC as shown *in vivo* (**Fig. 3H**). Increased levels of TFF1 and TFF2 in conjunction with mucus protein indicate the formation of a protective gastric mucous barrier in the hsMPS, which enables the study of pathogenesis within the *in vivo* relevant gastric mucous barrier.

### Enhancing the formation of epithelial barrier with gMSC co-culture under flow conditions

The integrity of the gastric epithelium is tightly regulated by functional apical-junctional complexes comprising tight and adherens junctions. Importantly, the adherens junction containing E-cadherin plays a key regulatory role in epithelial barrier integrity and highly associated with maturation and localization of secretory cells (*27*). To determine whether gMSCs contributes to the formation of junctional complexes in the hsMPS, we confirmed barrier integrity by observing E-cadherin and F-actin using confocal immunofluorescence assay. As shown in **Fig. 4A**, the integrity of E-cadherin in gastric epithelial cells was enhanced in co-culture conditions, which was again confirmed using qRT–PCR, presenting a seven-fold increase in mRNA expression of the CDH1 gene (**Fig. 4B**). Furthermore, F-actin was more concentrated in the cell-cell contact region presenting a higher co-localization efficiency of F-actin with E-cadherin (4.0±2.7% versus 15±2.2%) **(Figs. 4C and 4D**). E-cadherin stability depends on its linked to the actin cytoskeleton, thus, these data reveal that gMSCs facilitate the formation of structural integrity of the adherens junction. It is also noteworthy that gastric epithelial cells showed typical cobblestone-like phenotype of differentiated gastric epithelial cells when co-cultured with gMSCs because E-cadherin played a role in the maintenance of epithelial morphology and polarity (*28*). In contrast, in the absence of gMSCs, gastric epithelial cells showed spindle-like morphology (**Figs. 4A and 4E**), similar to a previous report (*5*). This demonstrates the role of gMSCs in recapitulating the gastric epithelial barrier and epithelial morphology in the hsMPS by enhancing the junctional complexes.

**Fig. 4.**
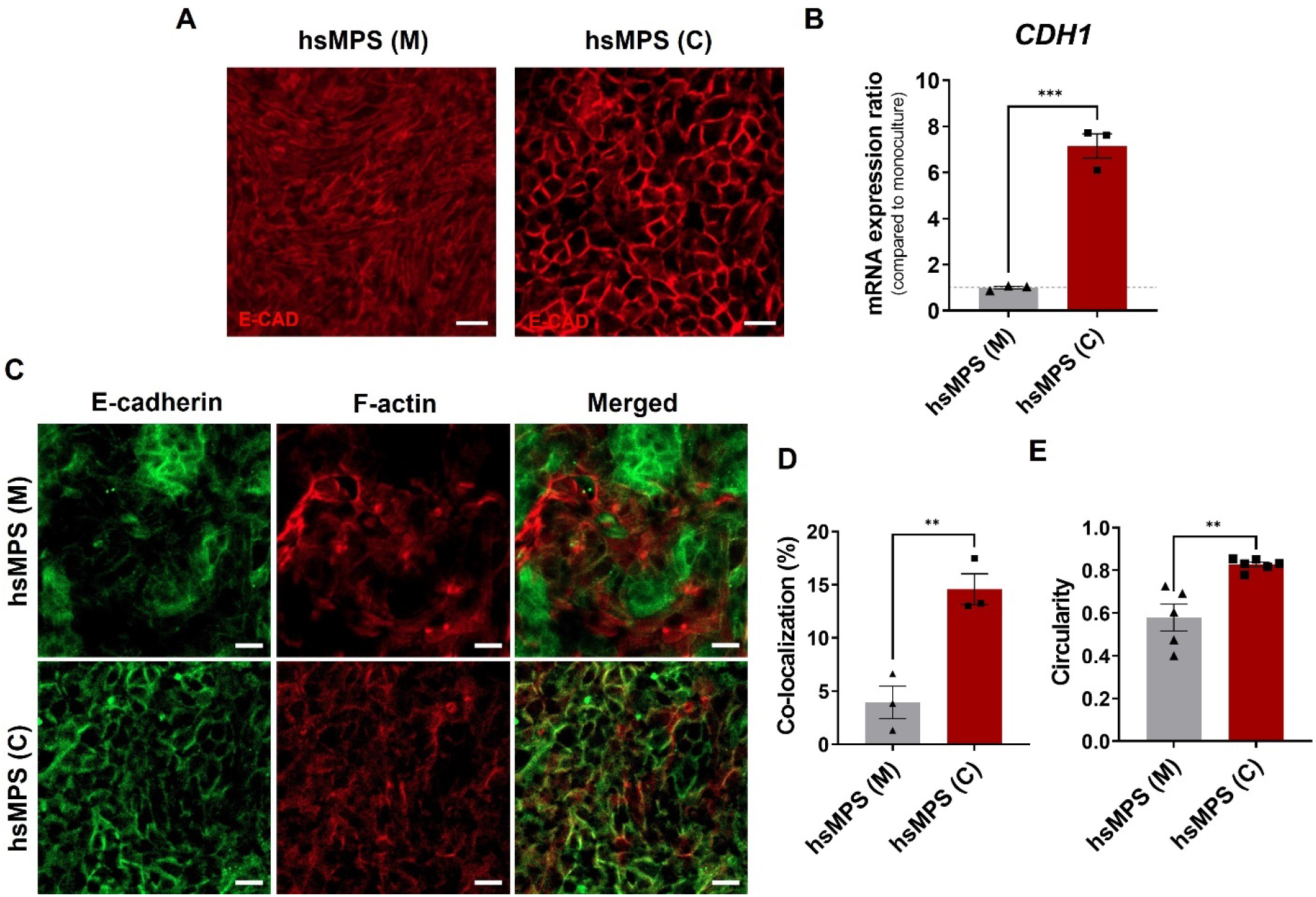
Epithelial junctional complexes in the hsMPS. (A) Immunofluorescence micrographic images of gastric epithelium in the mono-culture (M) and co-cultures (C) in the hsMPS labelled with E-cadherin (bar, 20 μm). (B) mRNA expression ratio of CDH1 encoding E-cadherin under co-culture conditions relative to monoculture conditions in the hsMPS. (C) Circularity of gastric epithelial cells in the monocultures and co-cultures analyzed using ImageJ. (D) Representative z-stacking immunofluorescence images of gastric epithelial cells in monoculture and co-culture conditions labelled with F-actin (red) and E-cadherin (green) (bar, 20 μm). (E) Co-localization efficiency of F-actin and E-cadherin analyzed using ImageJ. The results are presented as the mean ± s.e.m. For statistical analysis, an unpaired *t*-test was performed (\**P* < 0.05; \*\**P* < 0.01; \*\*\**P* < 0.001; \*\*\*\**P* < 0.0001).

### Use of hsMPS as the *H. pylori* infection model

We demonstrated that our hsMPS co-cultured with gMSCs under fluid flow exhibited characteristics of functional gastric epithelium with an *in vivo* relevant mucosal barrier. We then used the hsMPS to investigate the initial pathogenic process of *H. pylori* infection. The gastric epithelium was infected with CagA and VacA-positive strain J99 at various multiplicities of infection (MOIs), in the absence of media flow for 6 h, to allow *H. pylori* access to the gastric mucus layer. Subsequently, fresh DM was treated in laminar flow for another 18 h (**Fig. 5A**).

**Fig. 5.**
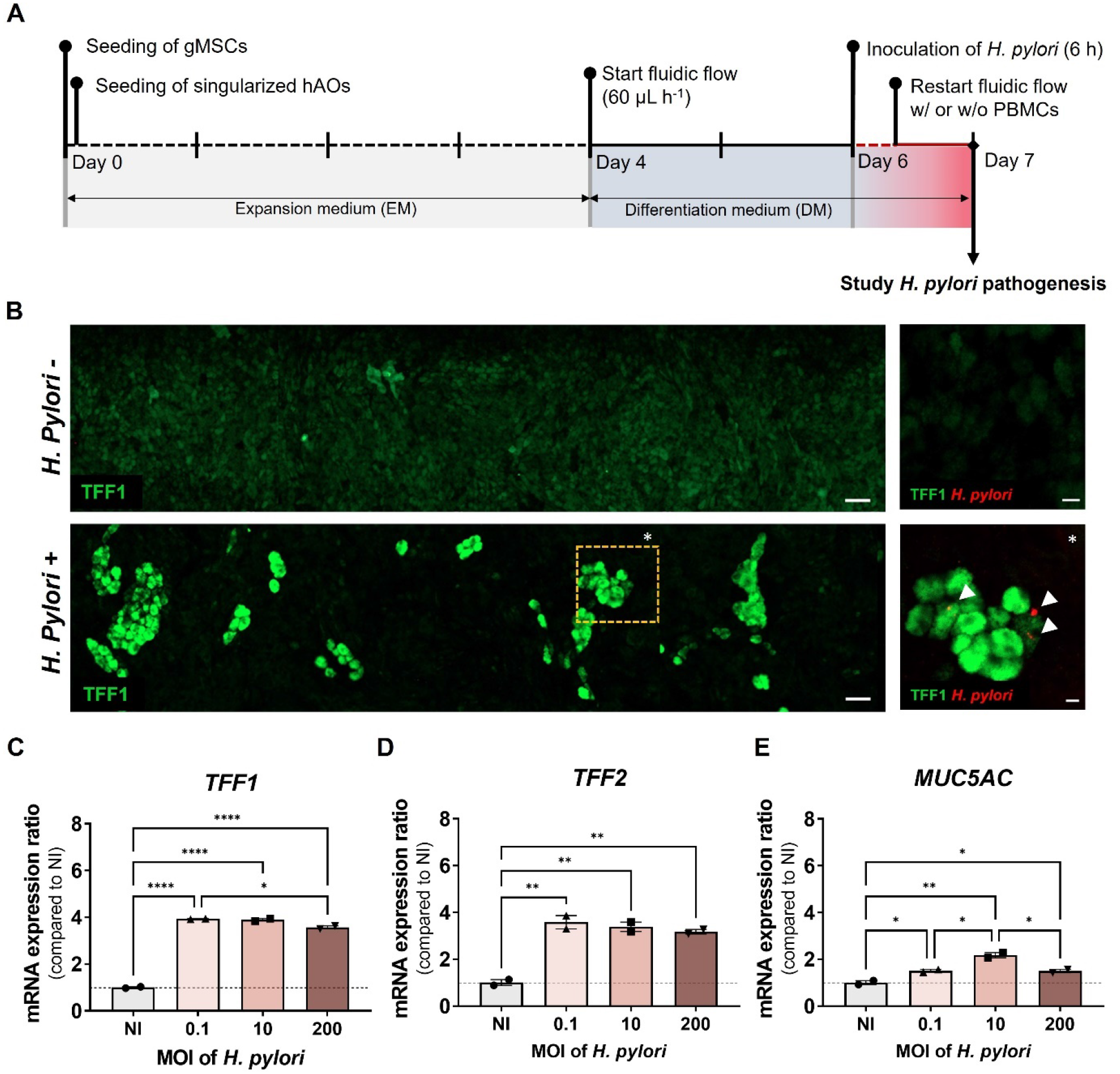
The protective function of gastric mucosal barrier against *H. pylori*infection. (A) Timeline for the generation of *H. pylori* infection model. At day 6, the gastric epithelium in the luminal microchannel of hsMPS was infected with *H. pylori* for 6 h in static condition and fresh DM was supplied at 60 μL h^−1^ for another 18 h. (B) Low magnification (left; bar, 50 μm) immunofluorescence micrographic images of the human gastric epithelium labelled with TFF1 (green) of a noninfected (top) and a *H. pylori*-infected hsMPS at an MOI 10 (bottom). High magnification (right; bar, 20 μm) immunofluorescence micrographic images showing gastric epithelium labelled with TFF1 (green) and *H. pylori* (red). (c-e) The mRNA expression ratios of TFF1 (C), TFF2 (D), and MUC5AC (E) in gastric epithelial cells when infected with *H. pylori* at various MOI relative to noninfected control. The results are presented as the mean ± s.e.m. For statistical analysis, one-way ANOVA and Tukey’s test was performed (\**P* < 0.05; \*\**P* < 0.01; \*\*\**P* < 0.001; \*\*\*\**P* < 0.0001). MOI, Multiplicity of infection. NI, non-infected.

Many studies have confirmed that TFF1 is upregulated in response to inflammation to protect the gastric epithelial secretions in autocrine and paracrine manners by performing anti-inflammatory functions (*29–31*). Thus, the pattern of TFF1 expression and localization of *H. pylori* was monitored via confocal immunofluorescence after the infection of the hsMPS with J99. Because histopathological studies indicate that MOI around 10 generally appears in *H. pylori* colonized gastric mucosa (*32*), an MOI of 10 was used to study the response of hsMPS to *H. pylori*. Interestingly, the expression of TFF1 was highly upregulated in distinct regions of the gastric epithelium in the vicinity of the *H. pylori* colonization zone, 24 h after infection (**Fig. 5B**). It is reasonable to assume that TFF1 islands were formed via paracrine action, inducing TFF1 expressions in nearby cells to intensively protect the epithelium from the spreading of *H. pylori*. Notably, the AGS cell line, which is known to have very low expression levels of TFF1 and MUC5AC, could not efficiently control the expansion of *H. pylori*, consequently leading to the coverage of the entire AGS epithelial layer by *H. pylori* in a MPS device (**fig. S3**).

We also analyzed the mRNA expression of genes encoding TFF1, TFF2, and MUC5AC after infection with *H. pylori* at different MOI values in the range from 0.1 to 200 (**Figs. 5C-5E**). Consistent with the confocal analysis, mRNA expression of TFF1 was significantly upregulated in *H. pylori*-positive hsMPS compared with that in the uninfected group, however, it was slightly lower at an MOI value of 200 compared with those at MOI values of 0.1 and 10. It was also revealed that mRNA expression levels of TFF2 and MUC5AC were significantly increased in *H. pylori-*positive sample, but not in MOI-dependent manner. This finding agrees with previous studies which revealed that TFF1, TFF2 and MUC5AC expressions are initially upregulated in acute *H. pylori* infection *in vivo*, while they are decreased in chronic infection (*33-35*), implying that our model reproduced the protective mechanism of the gastric mucosal barrier against *H. pylori* at an early stage. Additionally, the mRNA expression of gene encoding CDH1 was upregulated at MOI values in the range from 10 to 300 but decreased at MOI value of 600. This data suggested that the epithelial physical barrier was enhanced through the upregulation of cell-cell junctional molecule in acute *H. pylori* infection until certain value of MOI, but at higher MOI value, the protective mechanisms did not work effectively (**fig. S4**).

Finally, the features of the *H. pylori*-induced inflammatory immune responses orchestrated by sequential elaboration of proinflammatory cytokines were monitored in the hsMPS (**Fig. 6A-D**). We focused on the transcription factor nuclear factor kappa B (NF-κB) signaling pathway which is a known feature of *H. pylori* infections. Specifically, we analyzed the expression of genes encoding tumor necrosis factor alpha (TNF*a*), IL-8 (IL8), IL-1 beta (IL1B), lymphotoxin beta (LTB), and CC/CXC motif chemokine ligands (CXCL3, CXCL5, and CCL20), which were previously found to be overexpressed in patients infected with *H. pylori*. The mRNA expressions of IL1B and LTB genes as major NF-κB targets significantly increased at MOI of 10 and 200. TNFa and IL8, which are known to be upregulated through NF-κB pathway, also increased in MOI-dependent manner. Accordingly, the mRNA expression of multiple CC/CXC motif chemokine ligands (CXCL3, CXCL5, and CCL20) secreted from epithelial cells to recruit immune cells were upregulated in response to the NF-κB signaling (**Figs. 6E-6G**). No inflammatory response was observed when *H. pylori* infected at a MOI 0.1 and IL1B and LTB expressions in the hsMPS were rather downregulated compared with those in the non-infected group. This result is similar to that of a previous study that revealed that overexpression of TFF1 abrogated *H. pylori*-mediated inflammatory responses and resulted in the suppression of inflammatory and oncogenic activation (*36*). In this study, we observed that mRNA expressions of genes associated with gastric mucosal protective function (TFF1, TFF2, MUC5AC, and CDH1) were upregulated but did not increase depending on the infection dose. This might explain why inflammatory responses of gastric epithelial cells were sufficiently suppressed when they were infected with *H. pylori* at low MOI (0.1), but not controlled when infected at high MOI (10 and 200) in our model.

**Fig. 6.**
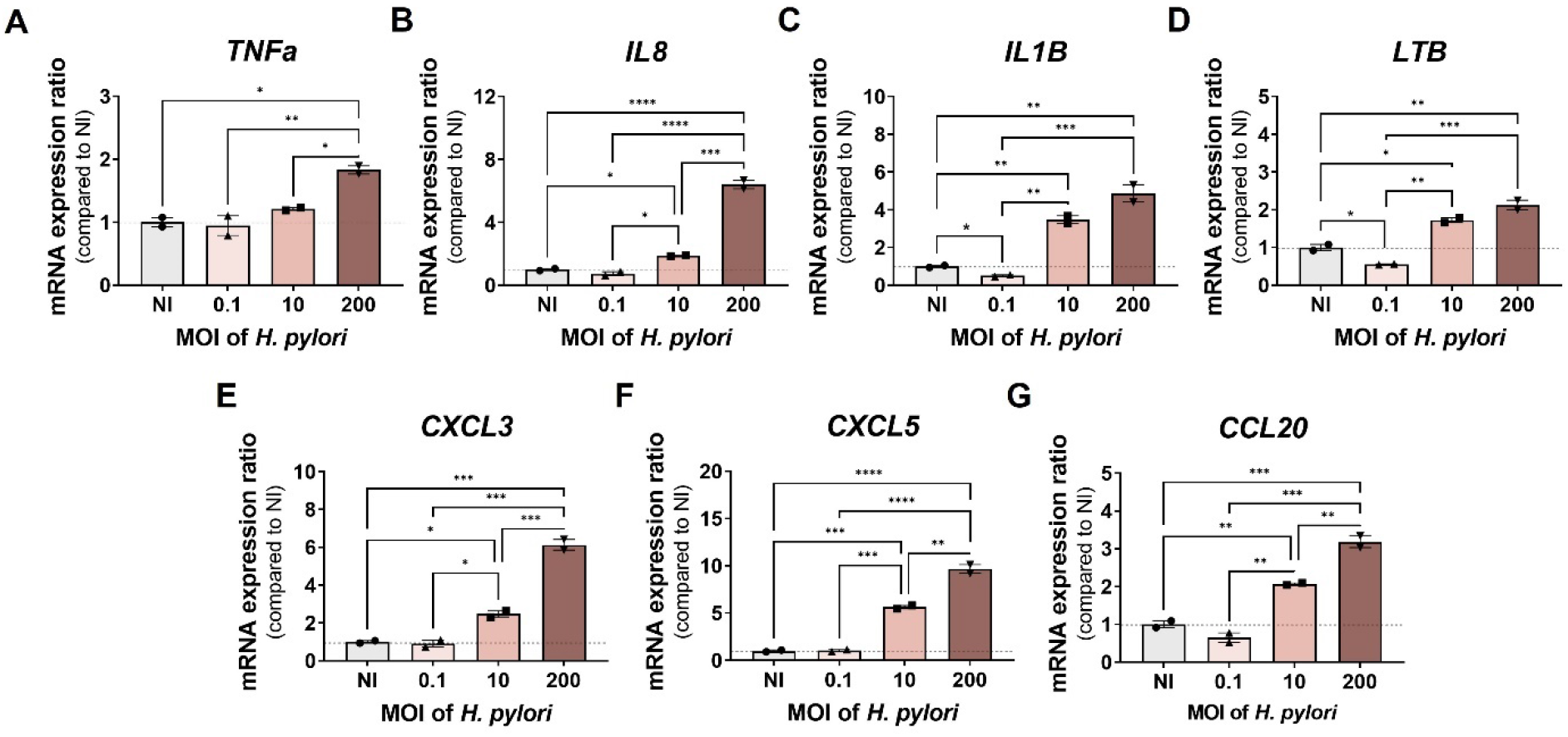
*H. pylori*-induced inflammatory responses in the hsMPS. The mRNA expression ratios of NF-κB-mediated inflammatory cytokines—TNFa (A), IL8 (B), IL1B (C), LTB (D), CXCL3 (E), CXCL5 (F), CCL 20 (G) in gastric epithelial cells when infected with *H. pylori* at various MOI relative to noninfected control. The results are presented as the mean ± s.e.m. For statistical analysis, one-way ANOVA and Tukey’s test was performed was performed. (\**P* < 0.05; \*\**P* < 0.01; \*\*\**P* < 0.001; \*\*\*\**P* < 0.0001). MOI, Multiplicity of infection. NI, non-infected. TNFa, tumor necrosis factor alpha; IL, interleukin; LTB, lymphotoxin beta; CCL/ CXCL, chemokine (C-C/C-X-C motif) ligand.

### An *in vitro* model to study immune pathogenic mechanism of *H. pylori* infection

As pro-inflammatory cytokines and chemokine ligands were increased in the hsMPS infected with *H. pylori* with activation of NF-κB signaling, we investigated whether our model could recapitulate the chemotaxis of immune cells shown in *H. pylori*-infected gastric tissue, which may lead to further chronic active inflammation in *H. pylori* infection (**Fig. 7A**). Human primary PBMCs were introduced in the abluminal channel, while *H. pylori* (MOI 10) was introduced into the luminal channel in the flow system (**Fig. 7A**). As expected, the number of PBMCs adhering to the abluminal side of the membrane was approximately 2.5-fold higher in the hsMPS infected with *H. pylori* than that in the non-infected MPS, emulating the immune cell recruitment at the *H. pylori* infection site (**Fig. 7B**). Because of the interplay between the gastric epithelium and immune cells, hsMPS system in combination with PBMC, showed a greatly enhanced level of responses in terms of the upregulation of NF-κB and cytokines expression (**Figs. 7C-7I**), as observed *in vivo*. In particular, IL-8 that is involved in all significant response pathways in the initial cellular response to *H. pylori* infection(*32*) (**Fig. 7E**), and CXCL3 (**Fig. 7G**) and CXCL5 (**Fig. 7H**), important neutrophil chemo-attractants (*37*) were highly activated in the presence of the immune cells, thus showing the contribution of immune cells in initial pathogenic mechanisms of *H. pylori* infection. These studies demonstrated the potential of the hsMPS as a promising *in vitro* tool to study immune pathogenic mechanism of *H. pylori* infection, and to further identify physiological contributions of individual immune cell types.

**Fig. 7.**
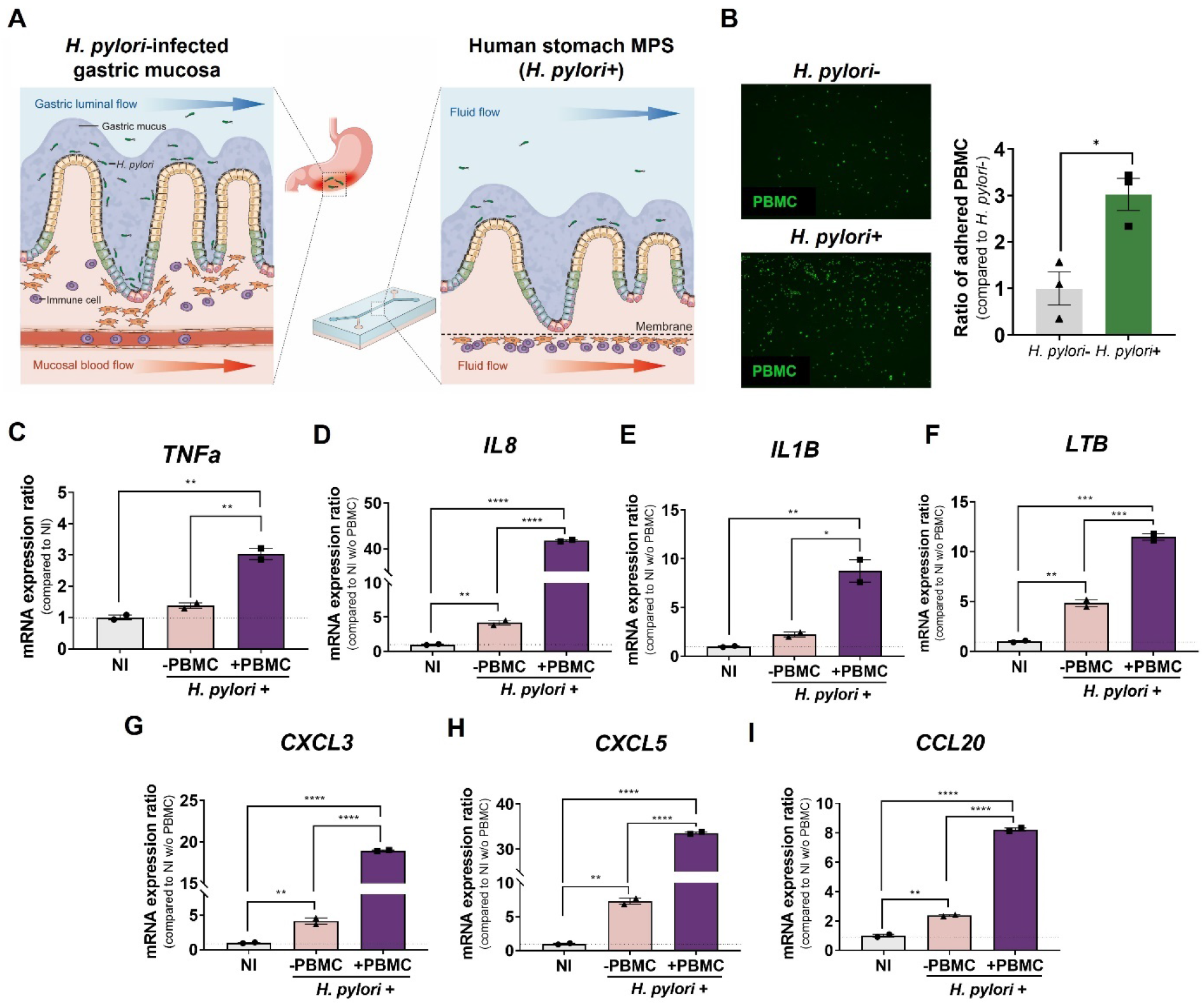
Immune pathogenesis in *H. pylori*-infected hsMPS co-cultivated with PBMC. (A) Schematic illustration of the *H. pylori*-infected human gastric mucosa (left) and *H. pylori*-infected hsMPS co-cultured with peripheral blood mononuclear cells (PBMCs). (B) Representative fluorescence images of PBMCs adhering on the basal side of the abluminal microchannel membrane of a non-infected (upper, left) and *H. pylori*-infected hsMPS at an MOI of 10 (lower, left) stained with CellTracker Green (bar, 250 μm). Relative number of PBMCs adhering on the basal side of the membrane of *H. pylori*-infected hsMPS at MOI 10 compared to non-infected control (right). (C-I) The mRNA expression ratios of NF-κB-mediated inflammatory cytokines— TNFa (C), IL8 (D), IL1B (E), LTB (F), CXCL3 (G), CXCL5 (H), CCL 20 (I) in gastric epithelial cells when infected with *H. pylori* at an MOI 10 in the presence and absence of PBMCs in abluminal microchannel relative to noninfected control (NI). The results are presented as the mean ± s.e.m. and n = 2 for the independent hsMPS experiments. For statistical analysis, one-way ANOVA with Tukey’s multiple comparisons test was performed (\**P* < 0.05; \*\**P* < 0.01; \*\*\**P* <0.001; \*\*\*\**P* < 0.0001). TNFa, tumor necrosis factor alpha; IL, interleukin; LTB, lymphotoxin beta; CCL/ CXCL, chemokine (C-C/C-X-C motif) ligand;-, without PBMC; +, with PBMC.

## Discussion

We report herein the first hsMPS, which demonstrated a significantly enhanced gastric mucosal barrier based on the combination of MPS technology with organoids. Our results demonstrated the formation of strong epithelial junctional complexes and an interconnected network of gastric mucus contained mucus-protecting peptides in the gastric epithelium in an MPS device. This was attributed to the recapitulation of homeostatic gastric stem cell renewal and differentiation induced by fluid flow and co-culture with gMSC. These features of the hsMPS enabled to model of gastric pathogen infections and related defense mechanisms by functional mucosal barrier, which is important for the development of effective therapeutic strategies.

Our study stems from previous studies using gastric organoids to generate a planar culture of the gastric epithelium in the Transwell systems (*5, 9*). Because gastric organoids provide gastric stem cells, which have the potential to proliferate or differentiate, gastric epithelium derived from organoids offers a relevant model by generating the cells required for creating components of the gastric mucosal barrier. Furthermore, the inoculation of *H. pylori* to the apical side of the epithelium can be performed more efficiently in planar cultures than in enclosed 3D organoids. The differentiation of gastric epithelial cells can be successfully induced by mimicking the gastric pit environment with attenuated Wnt signal. This was accomplished by restricting Wnt protein (Wnt3A) and its agonist (R-spondin1) in the culture media, while repressing gastric epithelial proliferation (*5, 9*). However, this approach does not fully reflect gastric epithelial physiology, which requires continuous renewal of the epithelium by gastric stem cells and generation of differentiation progeny. Thus, we employed MPS technology, which facilitates effective simulation of the *in vivo* microenvironment, including cell-cell interaction and perfusion, to recapitulate the dynamic renewal of gastric epithelial cells as well as cell differentiation observed in physiological conditions.

The mesenchymal components of gastric tissue have been suggested to regulate stem cell activity as a stem cell niche by secretion of mitogenic factors (*8*). It has already been demonstrated that gMSCs induce the proliferation of gastric stem cells using mouse gastric organoids and primary mouse gMSCs (*38*). Thus, we mimicked the human gastric stem cell niche by culturing gastric epithelial cells derived from hAOs interfaced with human gastric tissue-derived gMSCs in two-channel microfluidic chamber. We demonstrated the synergistic effect of co-cultures with gMSCs and fluid flow in renewal of gastric stem cells in differentiation media condition. The epithelial-mesenchymal interaction resulting in gastric epithelial homeostasis was only observed in the presence of perfusion, because potential of gMSCs as a cellular niche factor could be maintained via Notch activation under constant flow. It highlights the great advantage associated with the use of a microfluidic system. Under these conditions, the hsMPS exhibited more physiologically relevant levels of gastric stem cell markers (LGR5, CCK2R, AXIN2) compared to hAO culture. Due to the increase in the number of proliferating cells, buckling of the gastric epithelial layer observed in the development of gastric epithelium *in vivo* was also recapitulated, sharing a similarity with a previous finding using a microfluidic intestine model (*39*). We demonstrated that gMSCs cultured in flow conditions express higher level of paracrine factors including EGF and FGF10, however, we cannot rule out that other niche factors including other growth factors, hormone, and extracellular vesicles could also be involved in promotion of epithelial renewal. Considering that molecular mechanism of mesenchymal niche factor surrounding the gastric glands is little understood, further efforts are needed to characterize the interactions between gastric stem cells and gMSCs for better recapitulation of gastric epithelium homeostasis in the hsMPSs.

We also demonstrated that hsMPS provides a functionally mature gastric epithelium with a well-developed gastric mucosal barrier achieved by co-culturing with gMSCs and controlled fluid flow, which cannot be achieved in traditional organoid culture. Most importantly, secretion of mucin proteins found in the gastric antrum MUC5AC and MUC6 was stimulated when co-cultured with gMSCs at physiological level in the hsMPS. This resulted in the formation of *in vivo*-like interconnected mucus networks overlaid on the epithelium, and it provided physical protection against the pathogens as a gastric mucosal barrier. TFF peptides (TFF1 and TFF2) co-expressed with mucin proteins functioning as a chemical gastric mucosal barrier was also highly upregulated. Altogether, these data revealed how MPS technology potentiated the functional maturation of hAOs and resulted in the recapitulation of complex gastric mucosal barrier functions by recreating the microenvironment of homeostatic gastric epithelium. A previous study revealed that gastric epithelial cells showed effective mucin secretion when grown at the air-liquid interface (ALI); however, the effect of a TFF-mediated chemical barrier has not yet been well investigated (*9*). Adapting the ALI culture method to our hsMPS system might be an interesting research opportunity to enhance the function of the gastric mucosal barrier.

Furthermore, our gastric system demonstrated the formation of stronger epithelial adherens junctions, which was important for restricting paracellular transport and maintaining epithelial morphology and cell polarization. The formation of a mature adherens junction was examined by upregulating E-cadherin and bundled actin filaments connected to the cell junctions and by creating an epithelial physical barrier that limits the invasion of pathogens. Accompanied by enhanced epithelial junctional complexes, the epithelium appeared to have a cobblestone-like morphology as normal gastric epithelial cells, whereas epithelium cultured in the same MPS device without gMSCs had fibroblastic-like morphology consistent with a previous report(*5*).

In human gastric mucosa, *H. pylori* invasion of gastric epithelial cells is limited by the gastric mucus layer, which physically protects the epithelium, and TFF1 binds to *H. pylori* within gastric mucus for chemical protection. Due to the proinflammatory cytokines released from gastric epithelial cells, immune cells are recruited to the infected regions, and they initiate the mucosal immune responses. However, *H. pylori* has its own evasion strategies for the defense system of the host to survive and develop various gastric diseases(*40*). Hence, accurate approximation of gastric mucosal barrier functions provides a deep understanding of pathogen-host interactions and development of treatment methods. We recapitulated *H. pylori* pathogenesis and defense mechanisms *in vitro* using the hsMPS, which offers a physiologically relevant gastric mucosal barrier. The two-channel design of the hsMPS permits direct access to the luminal and abluminal compartments and facilitates the emulation of various infection conditions. MOI-dependent *H. pylori* pathogenesis was studied by adding *H. pylori* to luminal microchannel at MOI values of 0.1 to 200, and the immune cell responses were investigated by applying PBMCs into the abluminal microchannels. In our model, we confirmed that TFF peptides, E-cadherin, and mucin proteins were upregulated to defend against *H. pylori* infection not in a MOI-dependent manner. Upregulation of TFF1 appeared locally and formed TFF1 islands, which cannot be found using global gene expression analysis, and is associated with the chemical defense mechanism of the gastric mucosal barrier in efficiently blocking the spread of *H. pylori* within fenced regions. This finding might help to understand TFF1-mediated *H. pylori* defense mechanisms, which is less studied because of the lack of physiologically relevant *in vitro* model. The immune responses induced in gastric tissue infected with *H. pylori* were also evidenced in the hsMPS, including an increase in the canonical NF-kB pathway involved in immune responses and CC/CXC ligands in a MOI-dependent manner, as well as recruitment of immune cells. Therefore, the hsMPS provided a significant enhancement as it allowed for an improved understanding of gastric pathophysiology by recapitulating the *in vivo* environment of gastric mucosal barrier more effectively. Also, it may prove useful in the developing therapeutic strategies to control the pathological symptoms and in developing drugs that efficiently eradicate pathogens, as it provides an understanding of their evasion mechanism in overcoming the defense system of a host.

This study only focused on the recapitulation of the gastric mucosal barrier in the antrum region to study initial *H. pylori* infection and immune responses; however, gastric physiology and pathophysiology rely on the complex interplay between the antrum and corpus regions of the stomach. In the stomach, gastric acid, which determines the pH of gastric juice, is secreted from parietal cells in the corpus region (*41*). Gastric acid secretion in the corpus region is mediated by hormonal agents, including gastrin secreted from the antrum, as well as paracrine and neuronal pathways (*41*). Thus, a limitation of the current research is that intragastric pH was not appropriately recapitulated in our hsMPS, which only contained antrum stem cell-derived epithelial cells. The gastric pH can be also affected by pathogens such as *H. pylori* when spreading from the antrum mucosa to the corpus and inducing atrophy, hypergastrinemia, or gastric adenocarcinoma (*42*). Therefore, further work on developing MPS platforms which offer functionally linked antrum and corpus would help us study the underlying mechanisms of intragastric pH change under pathological conditions and investigate potential pharmacological approaches to gastric acid suppression for the treatment of acid-related disorders. Furthermore, hsMPS generated using patient-derived cells and pathogens will provide a key tool for precision medicine by predicting clinical complications depending on individual genetic and *H. pylori* virulence factors.

In summary, the hsMPS created using gastric epithelial cells derived from hAOs interfaced with gMSCs exhibited physiologically relevant gastric mucosal barrier functions. Gastric epithelial homeostasis and gastric mucosal barrier were successfully achieved by crosstalk between the epithelium and gMSCs, and mucosal flow in an MPS device. The functionalities provided enhanced epithelial junctional complexes creating epithelial barrier, *in vivo*-like continuous mucus barrier over the gastric epithelium with high levels of TFF1 and TFF2 expression and led to the recapitulation of infection-induced gastric epithelial responses. Therefore, our hsMPS provides significant advancement as the first stomach MPS that recapitulates the *in vivo* environment of the gastric mucosal barrier, it will improve drug development and therapeutic approaches for gastric bacterial and viral infection.

## Materials and Methods

### Generation of hAOs and gastric mesenchymal stromal cell culture

Human antrum was collected from patients who underwent surgery at the Seoul National University College of Medicine (IRB protocol number: H-2002-082-1102). The hAOs were generated as previously described (*43*) with some minor modifications. The antrum tissue was washed in Dulbecco's phosphate-buffered saline (DPBS) with antibiotics (Primocin and Plasmocin, InvivoGen) and was pinned on a silicone-coated Petri dish. The mucosa was carefully separated from the submucosa under a microscope using a microdissecting scissor and washed in ice-cold chelating buffer (DPBS without Ca^2+^/Mg^2+^ containing 1% sucrose, 2% D-sorbitol and 1% bovine serum albumin). The surface of the mucosa was scrapped gently using curved forceps to remove the gastric mucus. The mucosa was cut into small pieces (1–2 mm^2^) and incubated in 10 mM EDTA-chelating buffer on a shaker for 10 min at room temperature (RT). Tissue fragments were washed with ice-cold chelating buffer and covered with a glass slide on a petri dish. The gastric glands were isolated by applying pressure to the glass slide and directly suspended in advanced DMEM/F12 (Gibco) with 5% fetal bovine serum (FBS) in a conical tube. After centrifugation for 5 min at 250 g at 4℃, the pellet was resuspended in 20 μL Matrigel and cultured on a 48-well cell culture plate in the EM (**table S1**). The hAOs were passaged twice a week with a split ratio of 1:3 to 1:6.

The gMSC culture was established using gastric muscularis mucosae harvested from the gastric antrum tissue. The gastric muscularis mucosae was cut into small fragments (2–5 mm^2^) and washed in DPBS, and then placed in a buffer containing type Ⅰ collagenase (Sigma) and Accutase (Merck) for 1.5 h in a shaker at 37 ℃, after which the digest was treated with 5% FBS (Merck) to inactivate the enzymes. The dissociated cells and tissue fragments were centrifuged at 1,000 rpm for 5 mins at 4 ℃ and cultured in a T75 tissue-culture flask in a growth medium (advanced DMEM/F12 (Gibco) containing 5% bovine serum (Gibco)). The gMSCs were allowed to migrate from the tissue fragments attached to the dish and form a monolayer. The cells were cultured for approximately 7 days until they reached 80% confluency and then passaged for an establishment of a gMSC line.

### Device fabrication

The design of the hsMPS was modified from a previously reported human BBB-on-a-chip (*44*). The apical and basal channels were cast in polydimethylsiloxane (PDMS) (Sylgard 184) at a 10:1 ratio of base to curing agent in a custom acryl-patterned master mold. The microchannel was 2 cm long and 1 mm wide; top and bottom channels were 1 and 0.2 mm high, respectively. After curing the PDMS at 60 ℃ for 6 h, the PDMS replica was demolded from the master mold. Track-etched polyethylene terephthalate (PET) membranes (1 μm pore size) purchased from it4ip were bonded to PDMS as previously described. The PDMS parts and PET membranes were treated with oxygen plasma (80 sccm of O_2_ for 1 min). The PDMS parts were submerged in a 5% 3-aminopropyltriethoxysilane (Sigma) solution to generate surface amine, and PET membranes were submerged in a 1% 3-glycidoxypropyltriethoxysilane (Sigma) solution for epoxy functionalization for 20 min at RT. Parts were rinsed in water and dried with compressed air. The microfluidic device was assembled by placing a PET membrane between the PDMS parts and aligning them with the microchannel pattern. The assembled devices were placed in a 65 ℃ oven for 2 days to form a strong amine-epoxy bond.

### Reconstitution of hsMPS

The PDMS surface of the chip was activated by treating with plasma (30 sccm of air, for 1 min), and coated with 1% Matrigel for 1.5 h at 37 ℃. For co-culturing the gMSCs in a chip, gMSCs at a density of 1.25 × 10^6^ cells ml^−1^ were seeded on the abluminal channel in a gMSC medium. The chip was flipped immediately to allow the gMSCs to attach to the Matrigel-coated PET membrane, and then placed in the CO2 incubator. At 2 h after seeding gMSCs, the chip was flipped back, and the apical and basal channels were rinsed with EM. Gastric antrum epithelial cells were prepared by singularization of the hAOs by treatment with TrypLE express (Gibco) for 5 min at 37 ℃. Gastric antral epithelial cells at a density of 5 × 10^6^ cells ml^−1^ were seeded on the luminal channel in EM. After 24 h, the hsMPS was fed with fresh EM every day for additional three days. On the fourth day of cell seeding, the outlet channels of the chip were connected to syringe pumps (Fusion 200, Chemyx Inc.), and a DM (**table S1**) was flowed through the channels at 60 μL h^−1^. For static culture of hsMPS, the medium was switched to DM at day 4 and daily replaced with fresh DM to maintain the cells.

### Quantitative real-time PCR

The mRNA expression levels were analyzed using quantitative real-time polymerase chain reaction (qRT-PCR). Total RNA of cells within a microchannel was extracted using a RNeasy Kit (Qiagen) and cDNA was synthesized using a QuantiTech Reverse Transcription Kit (Qiagen) following the manufacturer's protocol. The qRT-PCR was carried out using a SYBR Green Realtime PCR Master Mix (TOYOBO) in a CFX Connect Real-Time PCR Detection System (Bio-Rad). The forward and reverse primers used for qRT-PCR are provided in **table S2**.

### Modeling *H. pylori* infection

*Helicobacter pylori* strain J99 (American Type Culture Collection 700824) was grown following the manufacturer’s protocol. Prior to infection into our chip, *H. pylori* was washed three times with DPBS and centrifuged for 3 min at 6000 × *g* at 4 ℃. *H. pylori* was gently inoculated into the luminal channel of the hsMPS and incubated in static condition. At 6 h after inoculation, the hsMPS was reconnected to the syringe pump (Fusion 200, Chemyx Inc.), and fresh culture media was provided to both microchannels at 60 μL h^−1^ for 24 h. To co-cultivate the *H. pylori* infection model with PBMCs, PBMCs were collected from peripheral blood of healthy donor (IRB protocol number: IRB-20-44-A) using Lymphoprep™ Density Gradient Medium (Stemcell Technologies Inc.) following the manufacturer’s protocol. Before adding PBMCs to the abluminal microchannel, we stained PBMCs using 5 μM of CellTracker Green (Invitrogen). The dyed PBMCs (1 × 10^6^ cells) were flowed into the abluminal channel after a 6-h inoculation. To observe the adhesion of *H. pylori* to the human gastric cell line, a stomach MPS that contains AGS cells (ATCC CRL-1739) was prepared using the same microfluidic device by culturing the AGS cells on the luminal channel and inoculated with *H. pylori* in the same manner as organoid-based hsMPS.

### Immunofluorescence microscopic analysis

The hsMPS was fixed with 4% paraformaldehyde in DPBS for 15 min and washed with DPBS 3 times for 5 min. Immunostaining was performed after permeabilization in DPBS with 0.1% Triton X-100 (Sigma) and blocking for 1 h in 10% goat serum in DPBS with 0.1% Triton X-100. The hsMPS microfluidic channels were treated with primary antibodies (**table S3**) and incubated overnight at 4 °C. After the washing procedure, secondary antibodies (**table S3**) were incubated for 1 h at RT. Nuclei were counterstained with 4′,6-diamidino-2-phenylindole (DAPI; Sigma). Conventional confocal imaging was carried out using confocal microscopes LSM780NLO (Zeiss) and LSM980 (Zeiss). To compare the gastric epithelial cell morphology between monoculture and co-culture, the area and perimeter of each cell were measured by using ImageJ software with immunofluorescence images of E-cadherin. Using each cell area and perimeter, the circularity could be calculated by the following formula:

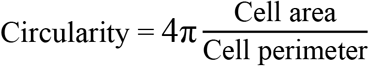

## Statistical analysis

All data represent means (±s.e.m). The statistical analysis was determined using a Student’s *t*-test to compare two sets of data, and one-way analysis of variance (ANOVA) with Tukey’s multiple comparisons test to compare in multiple groups; \**P* <0.05; \*\**P* <0.01; \*\*\**P* <0.001; \*\*\*\**P* < 0.0001). Prism 8 (GraphPad Software) was used for statistical analysis.

## Acknowledgments

We thank Hyun Myong Kim (Cancer Research Institute at Seoul National University) for assistance in preparing gastric tissue samples, Hye-Rim Shim (Ulsan National Institute of Science and Technology) for her excellent technical advice and providing *H. pylori* culture, and Seongjun You (Ulsan National Institute of Science and Technology) for assistance in image analysis.

## Funding

This study was supported by a National Research Foundation of Korea (NRF) grant funded by the Ministry of Science and ICT (NRF-2021R1A4A303059711 and NRF-2020R1C1C1014753), a Research Fund (1.210034.01) of Ulsan National Institute of Science and Technology, and a SNUH research fund (03-2020-0340) of Seoul National University Hospital.

## Author contributions

H.-J.J. participated in the design and performance of all experiments and analyzed the data with T.-E.P. and S.-H.K., who supervised all work. J.-H.P. and S.-H.K. obtained gastric tissue samples according to the IRB-approved protocol. J.K. provided scientific supervision on the experiments of *H. pylori* infection. H.-J.J. and T.-E.P. prepared the manuscript with input from all others.

## Competing interests

Authors declare that they have no competing interests.

## Data and materials availability

All data that support the findings of the study are included within the article and supplementary files.

## Notes

### Competing Interest Statement

The authors have declared no competing interest.

## References

1. L. E. Wroblewski, R. M. Peek Jr., Targeted disruption of the epithelial-barrier by Helicobacter pylori. Cell Commun. Signal. 9, 29 (2011).

2. S. Min, S. Kim, S.-W. Cho, Gastrointestinal tract modeling using organoids engineered with cellular and microbiota niches. Exp. Mol. Med. 52, 227–237 (2020).

3. S. Xiao, L. Zhou, Gastric Stem Cells: Physiological and Pathological Perspectives. Front. Cell. Dev. Biol. 8, 571536 (2020).

4. S. Bartfeld et al., In vitro expansion of human gastric epithelial stem cells and their responses to bacterial infection. Gastroenterol. 148, 126–136. e126 (2015).

5. P. Schlaermann et al., A novel human gastric primary cell culture system for modelling Helicobacter pylori infection in vitro. Gut 65, 202–213 (2016).

6. T. Sato et al., Long-term expansion of epithelial organoids from human colon, adenoma, adenocarcinoma, and Barrett’s epithelium. Gastroenterol. 141, 1762–1772 (2011).

7. T. L. Testerman, D. J. McGee, H. L. Mobley, Adherence and colonization. Helicobacter pylori physiology and genetics, 379–417 (2001).

8. T. Katano et al., Gastric mesenchymal myofibroblasts maintain stem cell activity and proliferation of murine gastric epithelium in vitro. Am. J. Pathol. 185, 798–807 (2015).

9. F. Boccellato et al., Polarised epithelial monolayers of the gastric mucosa reveal insights into mucosal homeostasis and defence against infection. Gut 68, 400–413 (2019).

10. T. A. Sebrell et al., A novel gastric spheroid co-culture model reveals chemokine-dependent recruitment of human dendritic cells to the gastric epithelium. Cell. Mol. Gastroenterol. Hepatol. 8, 157–171. e153 (2019).

11. S. E. Park, A. Georgescu, D. Huh, Organoids-on-a-chip. Science 364, 960–965 (2019).

12. Y. Wang, L. Wang, Y. Guo, Y. Zhu, J. Qin, Engineering stem cell-derived 3D brain organoids in a perfusable organ-on-a-chip system. RSC Adv. 8, 1677–1685 (2018).

13. M. Kasendra et al., Development of a primary human Small Intestine-on-a-Chip using biopsy-derived organoids. Sci. Rep. 8, 2871 (2018).

14. K. Achberger et al., Merging organoid and organ-on-a-chip technology to generate complex multi-layer tissue models in a human retina-on-a-chip platform. Elife 8, (2019).

15. K. A. Homan et al., Flow-enhanced vascularization and maturation of kidney organoids in vitro. Nat. Methods 16, 255–262 (2019).

16. K. K. Lee et al., Human stomach-on-a-chip with luminal flow and peristaltic-like motility. Lab Chip 18, 3079–3085 (2018).

17. H. J. Kim, D. Huh, G. Hamilton, D. E. Ingber, Human gut-on-a-chip inhabited by microbial flora that experiences intestinal peristalsis-like motions and flow. Lab Chip 12, 2165–2174 (2012).

18. S. Kawano, S. Tsuji, Role of mucosal blood flow: a conceptional review in gastric mucosal injury and protection. J. Gastroenterol. Hepatol. 15, 1–6 (2000).

19. M. Sigal et al., Stromal R-spondin orchestrates gastric epithelial stem cells and gland homeostasis. Nature 548, 451–455 (2017).

20. Y. Inoue, T. Watanabe, S. Okuda, T. Adachi, Mechanical role of the spatial patterns of contractile cells in invagination of growing epithelial tissue. Dev. Growth. Differ. 59, 444–454 (2017).

21. Y. Tian et al., Notch activation enhances mesenchymal stem cell sheet osteogenic potential by inhibition of cellular senescence. Cell Death Dis 8, e2595 (2017).

22. M. Clyne et al., Helicobacter pylori interacts with the human single-domain trefoil protein TFF1. Proc. Natl. Acad. Sci. U. S. A. 101, 7409–7414 (2004).

23. M. Clyne, F. E. May, The Interaction of Helicobacter pylori with TFF1 and Its Role in Mediating the Tropism of the Bacteria within the Stomach. Int. J. Mol. Sci. 20, 4400 (2019).

24. A. J. Peterson et al., Helicobacter pylori infection promotes methylation and silencing of trefoil factor 2, leading to gastric tumor development in mice and humans. Gastroenterology 139, 2005–2017 (2010).

25. W. Liu et al., Trefoil factor 1 and gastrokine 2 inhibit Helicobacter pylori-induced proliferation and inflammation in gastric cardia and distal carcinogenesis. Oncol Lett 20, 318 (2020).

26. X. Murgia, B. Loretz, O. Hartwig, M. Hittinger, C. M. Lehr, The role of mucus on drug transport and its potential to affect therapeutic outcomes. Adv Drug Deliv Rev 124, 82–97 (2018).

27. M. R. Schneider et al., A key role for E-cadherin in intestinal homeostasis and Paneth cell maturation. PLoS One 5, e14325 (2010).

28. M. Fujiwara et al., Epithelial DLD-1 Cells with Disrupted E-cadherin Gene Retain the Ability to Form Cell Junctions and Apico-basal Polarity. Cell Struct. Funct. 40, 79–94 (2015).

29. P. Thuwajit et al., Increased TFF1 trefoil protein expression in Opisthorchis viverrini‐ associated cholangiocarcinoma is important for invasive promotion. Hepatol. Res. 37, 295–304 (2007).

30. N. Amiry et al., Trefoil factor-1 (TFF1) enhances oncogenicity of mammary carcinoma cells. Endocrinology 150, 4473–4483 (2009).

31. D. R. Radiloff et al., Trefoil factor 1 acts to suppress senescence induced by oncogene activation during the cellular transformation process. Proc. Natl. Acad. Sci. U. S. A. 108, 6591–6596 (2011).

32. L. L. Eftang, Y. Esbensen, T. M. Tannaes, I. R. Bukholm, G. Bukholm, Interleukin-8 is the single most up-regulated gene in whole genome profiling of H. pylori exposed gastric epithelial cells. BMC Microbiol. 12, 9 (2012).

33. R. Esposito et al., Gastric TFF1 Expression from Acute to Chronic Helicobacter Infection. Front. Cell Infect. Microbiol. 7, 434 (2017).

34. W. Gonciarz et al., Upregulation of MUC5AC production and deposition of LEWIS determinants by HELICOBACTER PYLORI facilitate gastric tissue colonization and the maintenance of infection. J. Biomed. Sci. 26, 23 (2019).

35. G. Y. Hu et al., Expression of TFF2 and Helicobacter pylori infection in carcinogenesis of gastric mucosa. World J. Gastroenterol. 9, 910–914 (2003).

36. M. Soutto et al., NF-kB-Dependent Activation of STAT3 by H. Pylori is Suppressed by TFF1. Res. Sq., (2021).

37. Z. Mao et al., CXCL5 promotes gastric cancer metastasis by inducing epithelial-mesenchymal transition and activating neutrophils. Oncogenesis 9, 63 (2020).

38. M. A. Schumacher et al., The use of murine-derived fundic organoids in studies of gastric physiology. J. Physiol. 593, 1809–1827 (2015).

39. W. Shin, C. D. Hinojosa, D. E. Ingber, H. J. Kim, Human Intestinal Morphogenesis Controlled by Transepithelial Morphogen Gradient and Flow-Dependent Physical Cues in a Microengineered Gut-on-a-Chip. iScience 15, 391–406 (2019).

40. A. Karkhah et al., Helicobacter pylori evasion strategies of the host innate and adaptive immune responses to survive and develop gastrointestinal diseases. Microbiol. Res. 218, 49–57 (2019).

41. A. C. Engevik, I. Kaji, J. R. Goldenring, The Physiology of the Gastric Parietal Cell. Physiol. Rev. 100, 573–602 (2020).

42. J. G. Kusters, A. H. van Vliet, E. J. Kuipers, Pathogenesis of Helicobacter pylori infection. Clin. Microbiol. Rev. 19, 449–490 (2006).

43. S. Bartfeld, H. Clevers, Organoids as model for infectious diseases: culture of human and murine stomach organoids and microinjection of Helicobacter pylori. J. Vis. Exp., (2015).

44. T. E. Park et al., Hypoxia-enhanced Blood-Brain Barrier Chip recapitulates human barrier function and shuttling of drugs and antibodies. Nat. Commun. 10, 2621 (2019).

