## Supplementary Fig 1 for "Organoid-based human stomach micro-physiological system to recapitulate the dynamic mucosal defense mechanism"

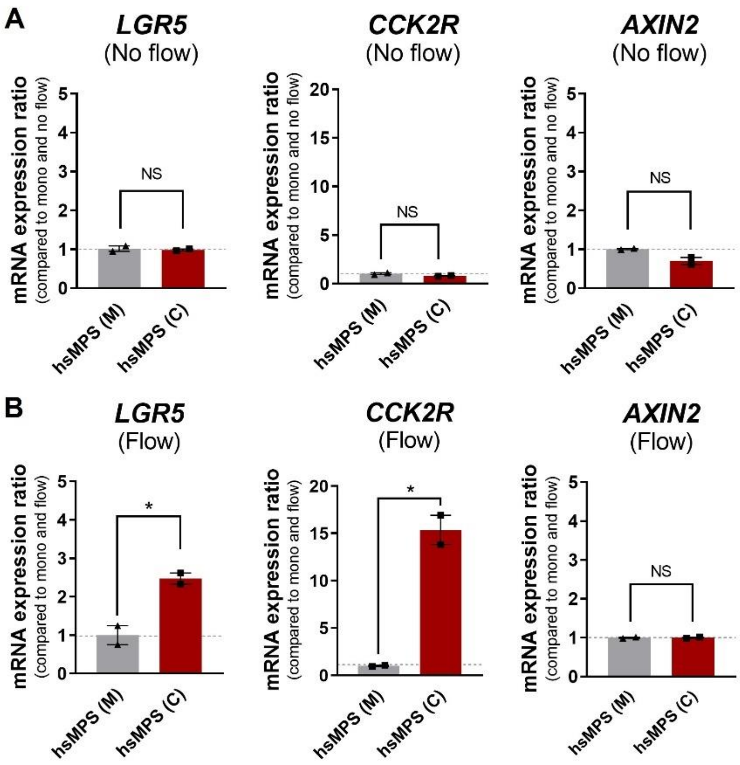


**fig. S1. Fluid flow-enhanced gastric stemness in the hsMPS co-cultured with gMSCs.** The mRNA expression ratio of LGR5, CCK2R, and AXIN2 in the (A) static (No flow) and (B) flow condition. The results are presented as the mean ± s.e.m. For statistical analysis, one-way ANOVA with Tukey's multiple comparisons test was performed (*$P < 0.05$; **$P < 0.001$; ***$P <0.001$; ****$P < 0.0001$).
