## Supplementary Fig. 2 for "Organoid-based human stomach micro-physiological system to recapitulate the dynamic mucosal defense mechanism"

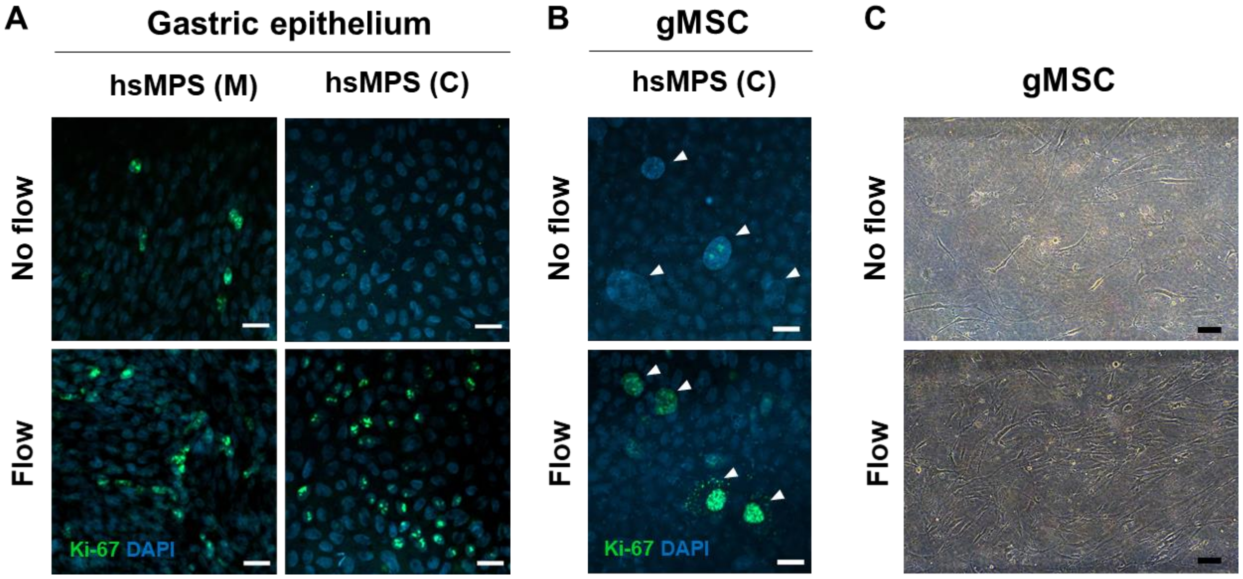


**fig. S2. Mitotic activity of gastric epithelium and mesenchymal stromal cells (gMSCs) in the hsMPS.** (A) Immunofluorescence images of Ki-67-positive gastric epithelial cells to compare mono-cultures (M) and co-cultures (C), and static (No flow) and flow condition in the hsMPS. (B) Immunofluorescence images of Ki-67-positive gMSCs in static (top) and flow condition (bottom). The nucleus of gMSCs was indicated by the white arrow. (C) Brightfield images showing density of gMSCs in static (top) and flow condition (bottom) in the hsMPS. White and black bar are indicated by 20 μm and 100 μm, respectively.
