## Supplementary Fig. 3 for "Organoid-based human stomach micro-physiological system to recapitulate the dynamic mucosal defense mechanism"

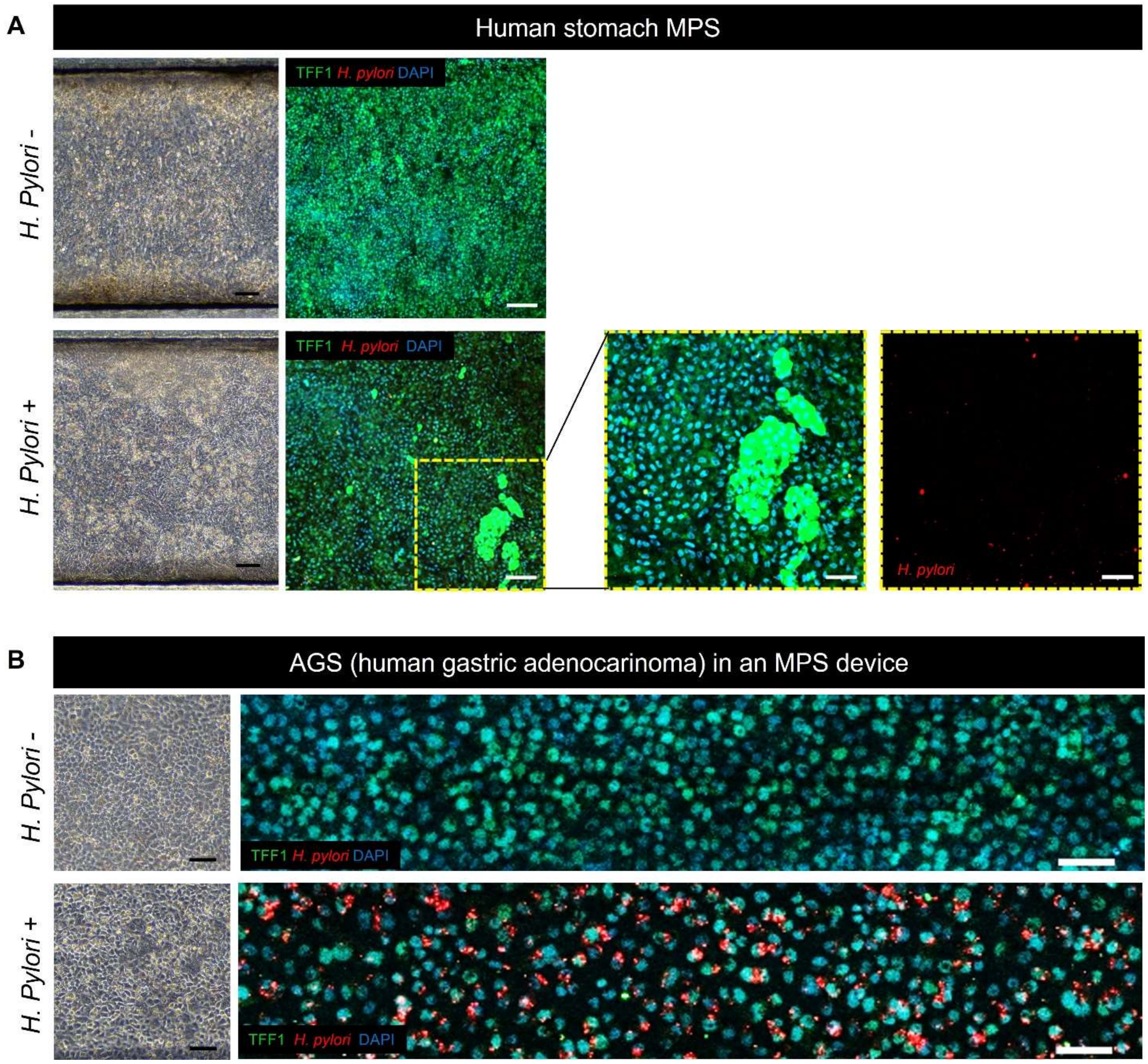


**fig. S3. *H. pylori*-infected gastric epithelium labeled with TFF1 and *H. pylori* in the hsMPS and AGS cells cultured in an MPS device.** (A) Brightfield microscopic images of non-infected (top, left) and *H. pylori*-infected (bottom, left) hsMPS. Low magnification immunofluorescence image of hsMPS of non-infected (top right) and *H. pylori*-infected at a MOI 10 (bottom, center) labeled with TFF1 (green), *H. pylori* (red), and nucleus (DAPI, blue) (bar, 100 μm). High magnification immunofluorescence images marked by yellow dotted box showing TFF1 island and *H. pylori* (bottom, right) (bar, 50 μm). (B) Brightfield microscopic images of AGS cells of non-infected (top, left) and *H. pylori*-infected at a MOI 10 (bottom, left) cultured in an MPS device (bar, 100 μm). The TFF1 expression level is shown decreased in *H. pylori*-infected AGS cells compared to that of non-infected cells. Fluorescence intensity of labeled *H. pylori* was relatively higher in AGS cells compared to that of hsMPS (bar, 50 μm). MOI, Multiplicity of infection.
