## Supplementary Fig. 4 for "Organoid-based human stomach micro-physiological system to recapitulate the dynamic mucosal defense mechanism"

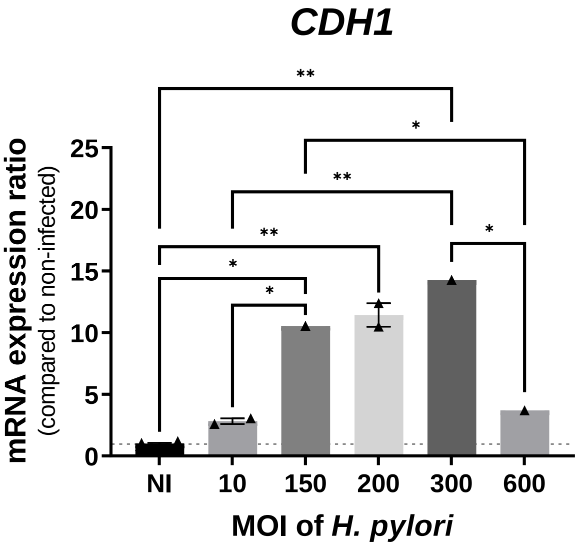


**fig. S4. The mRNA expression ratio of CDH1 in gastric epithelial cells when infected with *H. pylori* at various MOIs relative to non-infected (NI) group.** The results are presented as the mean ± s.e.m. For statistical analysis, one-way ANOVA with Tukey's multiple comparisons test was performed (**P* < 0.05; ***P* < 0.001; ****P* <0.001; *****P* < 0.0001).
