## Supplementary Table 1 for "Organoid-based human stomach micro-physiological system to recapitulate the dynamic mucosal defense mechanism"

**table S1. Composition of expansion media (EM) and differentiation media (DM)**

| No. | Component | Manufacturer | Catalog no. | EM | DM |
|---|---|---|---|---|---|
| 1 | Advanced DMEM/F12 | Gibco | 12634028 | O | O |
| 2 | L-WRN conditioned media (50%) | ATCC | CRL-3276 | O | - |
| 3 | Noggin (100 ng/ml) | Peprotech | 200-10C | - | O |
| 4 | Primocin (100 μg/ml) | InvivoGen | ant-pm-1 | O | O |
| 5 | Plasmocin (5 μg/ml) | InvivoGen | ant-mpp | O | O |
| 6 | GlutaMAX (2 mM) | Gibco | 35050061 | O | O |
| 7 | HEPES (10 mM) | Welgene | 15630106 | O | O |
| 8 | Nicotinamide (10 mM) | Sigma | N3376 | O | O |
| 9 | N-acetyl-L-cysteine (1.25 mM) | Sigma | A9165 | O | O |
| 10 | B27 supplement (1X) | Gibco | 17504044 | O | O |
| 11 | Recombinant Human FGF10 (20 ng/ml) | Peprotech | 100-26 | O | O |
| 12 | Animal-Free Recombinant Human EGF (50 ng/ml) | Peprotech | AF-100-15 | O | O |
| 13 | A83-01 (500 nM) | TOCRIS | 2939 | O | O |
| 14 | Gastrin - [Leu15]-Gastrin I human | Sigma | G9145 | O | O |
| 15 | Y27632 (10 μM) | TOCRIS | 1254 | O | O |
| 16 | Fetal bovine serum (5%) | Merck | TMS-013-BKR | - | O |
