## Supplementary Table 2 for "Organoid-based human stomach micro-physiological system to recapitulate the dynamic mucosal defense mechanism"

### table S2. List of primer sequences used for qRT-PCR

| | | | | | |
|---|---|---|---|---|---|
| **MUC5AC** | Forward | CAGCACAACCCCTGTTTCAAA | **NOTCH1** | Forward | TCAGCGGGATCCACTGTGAG |
| | Reverse | GCGCACAGAGGATGACAGT | | Reverse | ACACAGGCAGGTGAACGACTTG |
| **MUC6** | Forward | CTGCCCTATACCAGCAATGGA | **NOTCH2** | Forward | ACAGTTGTGTCTGCTCACCAGGAT |
| | Reverse | CTGACCCATGTACTTCCGCTC | | Reverse | GCGGAAACCATTCACACCGTTGAT |
| **GAST** | Forward | ATGCAGCGACTATGTGTGTATG | **NOTCH3** | Forward | GATGGCATGGATGTCAATGTTCGT |
| | Reverse | GCCCCTGTACCTAAGGGTG | | Reverse | TGCCTCATCCTCTTCAGTTGGCAT |
| **SST** | Forward | ACCCAACCAGACGGAGAATGA | **NOTCH4** | Forward | TCAAACAGAGGTGGATGAGTGCCT |
| | Reverse | GCCGGGTTTGAGTTAGCAGA | | Reverse | AGTTGGCCTTGTCTTTCTGGTCCT |
| **TFF1** | Forward | CCCTCCCAGTGTGCAAATAAG | **DLL1** | Forward | TGTGACGAGTGTATCCGCTATCCA |
| | Reverse | GAACGGTGTCGTCGAAACAG | | Reverse | AGGGCTTATGGTGTGTGCAGTAGT |
| **TFF2** | Forward | CGGGGAGTGAGAAACCCTC | **DLL3** | Forward | TCCCGGATGCACTCAACAACCTAA |
| | Reverse | CACTGGAGTCGAAACAGCATC | | Reverse | TTCAGGGCGATTCCAATCTACGGA |
| **CDH1** | Forward | ATTTTTCCCTCGACACCCGAT | **DLL4** | Forward | ACTGCGAGAAGAAAGTGGACAGGT |
| | Reverse | TCCCAGGCGTAGACCAAGA | | Reverse | ACATGAGCCCATTCTCCAGGTCAT |
| **TNFα** | Forward | TCCCCAGGGACCTCTCTCTA | **JAG1** | Forward | ACTGCTCACCCTGAAAGACCACT |
| | Reverse | GAGGGTTTGCTACAACATGGG | | Reverse | AGGACCACAGACGTTCCAGGAAAT |
| **IL-8** | Forward | ACACTGCGCCAACACAGAAAT | **JAG2** | Forward | TGCTGTGGAGGTGGCTATGTCT |
| | Reverse | ATTGCATCTGGCAACCCTACA | | Reverse | TGTTTCCACCTTGACCTCGGT |
| **IL-1β** | Forward | AGCTACGAATCTCCGACCAC | **HES1** | Forward | GTCAACACGACACCGGATAAACCA |
| | Reverse | CGTTATCCCATGTGTCGAAGAA | | Reverse | TTTCCAGAATGTCCGCCTTCTCCA |
| **CXCL1-3** | Forward | CGCCCAAACCGAAGTCATAG | **LGR5** | Forward | GAGTTACGTCTTGCGGGAAAC |
| | Reverse | GCTCCCCTTGTTCAGTATCTTTT | | Reverse | TGGGTACGTGTCTTAGCTGATTA |
| **CXCL5** | Forward | GATCCAGAAGCCCCTTTTCT | **CCK2R** | Forward | CGTGTGCTGCAGTGCGTGCA |
| | Reverse | GAAACTTTTCCATGCGTGCT | | Reverse | GGTGGTGTAGCTAAGCCTGG |
| **CCL20** | Forward | TGCTGTACCAAGAGTTTGCTC | **AXIN2** | Forward | TACACTCCTTATTGGGCGATCA |
| | Reverse | CGCACACAGACAACTTTTTCTTT | | Reverse | TTGGCTACTCGTAAAGTTTTGGT |
| **LTB** | Forward | GTACGGGCCTCTCTGGTACA | **EGF** | Forward | TGTCCACGCAATGTGTCTGAA |
| | Reverse | GTCCACCATATCGGGGTGAC | | Reverse | CATTATCGGGTGAGGAACAACC |
| **GAPDH** | Forward | TGTGGGCATCAATGGATTTGG | **FGF10** | Forward | CAGTAGAAATCGGAGTTGTTGCC |
| | Reverse | ACACCATGTATTCCGGGTCAAT | | Reverse | TGAGCCATAGAGTTTCCCCTTC |
| | | | **RSPO3** | Forward | TGTGCAACATGCTCAGATTACA |
| | | | | Reverse | TGCTTCATGCCAATTCTTTCCA |
