## Supplementary Table 3 for "Organoid-based human stomach micro-physiological system to recapitulate the dynamic mucosal defense mechanism"

### table S3. List of antibodies used for immunofluorescence micrographic analysis

| Primary antibody | Manufacturer | Catalog no. | Target name | Host |
|---|---|---|---|---|
| 1 | Invitrogen | MA5-12178 | Mucin 5AC | Mouse |
| 2 | Santa cruz | sc-33668 | Mucin 6 | Mouse |
| 3 | Santa cruz | sc-28302 | Gastrin | Mouse |
| 4 | Invitrogen | PA5-87974 | LGR5 | Rabbit |
| 5 | Proteintech | 20874-1-AP | E-cadherin | Rabbit |
| 6 | Santa cruz | sc-23900 | Ki67 | Mouse |
| 7 | Proteintech | 13734-1-AP | TFF1 | Rabbit |
| 8 | Abcam | ab268118 | TFF1 | Mouse |
| 9 | Sigma | A22287 | Phalloidin | - |
| 10 | Abcam | ab20459 | *H. pylori* | Rabbit |

| Secondary antibody | Manufacturer | Catalog no. | Product name | Host |
|---|---|---|---|---|
| 1 | Invitrogen | A11001 | Goat anti-Mouse IgG (H+L) Cross-Adsorbed Secondary Antibody, Alexa Fluor 488 | Goat |
| 2 | Invitrogen | A110034 | Goat anti-Rabbit IgG (H+L) Highly Cross-Adsorbed Secondary Antibody, Alexa Fluor 488 | Goat |
| 3 | Abcam | ab150083 | Goat Anti-Rabbit IgG H&L (Alexa Fluor® 647) | Goat |
